# Riboswitch and small RNA modulate *btuB* translation initiation in *Escherichia coli* and trigger distinct mRNA regulatory mechanisms

**DOI:** 10.1101/2023.11.17.567546

**Authors:** L. Bastet, A.P. Korepanov, J. Jagodnik, J.P. Grondin, A.M. Lamontagne, M. Guillier, D.A. Lafontaine

## Abstract

Small RNAs (sRNAs) and riboswitches represent distinct classes of RNA regulators that control gene expression upon sensing metabolic or environmental variations. While sRNAs and riboswitches regulate gene expression by affecting mRNA and protein levels, existing studies have been limited to the characterization of each regulatory system in isolation, suggesting that sRNAs and riboswitches target distinct mRNA populations. We report that the expression of *btuB* in *Escherichia coli*, which is regulated by an adenosylcobalamin (AdoCbl) riboswitch, is also controlled by the small RNAs OmrA and, to a lesser extent, OmrB. Strikingly, we find that the riboswitch and sRNAs reduce mRNA levels through distinct pathways. Our data show that while the riboswitch triggers Rho-dependent transcription termination, sRNAs rely on the degradosome to modulate mRNA levels. Importantly, OmrA pairs with the *btuB* mRNA through its central region, which is not conserved in OmrB, indicating that these two sRNAs may have specific targets in addition to their common regulon. In contrast to canonical sRNA regulation, we find that OmrA repression of *btuB* is lost using an mRNA binding-deficient Hfq variant. Together, our study demonstrates that riboswitch and sRNAs modulate *btuB* expression, providing an example of *cis-* and *trans-*acting RNA-based regulatory systems maintaining cellular homeostasis.

## Introduction

Bacteria adapt to environmental changes by modulating gene expression at the mRNA and protein levels (1, 2). Non-coding RNAs are involved in multiple bacterial regulatory processes and have been shown to be crucial actors in adaptive responses (2). Among these, riboswitches are RNA regulators often located in the 5’ untranslated region (5’ UTR) that modulate gene expression by undergoing structural changes (3, 4). These metabolite-binding RNA regulators control gene expression at the levels of transcription termination, translation initiation or mRNA decay (3). In several cases, metabolite binding to riboswitches has been shown to affect multiple regulatory processes, suggesting that these RNA elements orchestrate complex regulatory pathways. For example, lysine binding to the *lysC* riboswitch downregulates gene expression both by directing RNase E cleavage of *lysC* mRNA and by inhibiting translation initiation (5). Furthermore, the *lysC* riboswitch was recently proposed to modulate Rho-dependent transcription termination (6, 7), indicating that the lysine riboswitch may control at least three regulatory processes to ensure tight genetic regulation. Another example of multiple mechanisms involved in genetic regulation was also characterized in *Corynebacterium glutamicum* where the flavin mononucleotide (FMN) riboswitch controls both RNase E/G cleavage activity and Rho-dependent transcription termination (8). Although such complex systems involving multiple regulatory factors are expected to be largely used by bacteria, relatively few studies address how they are coordinated during the regulation process, i.e., whether regulatory activities are performed independently or if they require any hierarchical order to achieve a timely and appropriate regulation.

Small RNAs (sRNAs) are regulatory elements that are often involved in cellular adaptative responses (2). In contrast to riboswitches, sRNAs typically control gene expression through intermolecular base pairing with targeted mRNAs and may involve an RNA chaperone, such as ProQ or the Sm-like protein Hfq, to facilitate mRNA recognition (9–12). sRNAs control transcription termination, translation initiation and mRNA degradation by recognizing sequences located in untranslated or coding regions (2). Because sRNAs often bind mRNA sequences using limited base pairing complementarity, this allows for a given sRNA to regulate multiple mRNA targets, thereby enabling a global regulatory response. Surprisingly, although both riboswitches and sRNAs mostly modulate gene expression by targeting mRNA untranslated regions, evidence for both effectors controlling the expression of the same mRNAs is lacking, entailing that sRNAs and riboswitch regulatory mechanisms may be mutually exclusive and that they might target globally different mRNA populations.

The *E. coli btuB* riboswitch regulates the synthesis of the BtuB membrane transporter that mediates the influx/efflux of corrinoids such as vitamin B_12_ (13, 14). Upon binding to adenosylcobalamin (AdoCbl)—one of the active forms of vitamin B_12_—the AdoCbl riboswitch prevents *btuB* translation initiation by sequestering the Shine-Dalgarno (SD) sequence in a stem-loop structure (Figure 1A)(15–18). Previous microarray data (19) suggested that *btuB* expression is also repressed by two highly similar sRNAs, OmrA and OmrB, whose genes are located adjacently to each other within the *aas* and *galR* intergenic region in *E. coli* (Figure 1A) (19). These sRNAs share nearly identical 5’ and 3’ ends (Supplementary Figure S1A) and are predicted to exhibit specific secondary structures (Supplementary Figure S1B). They are negative regulators of genes encoding outer membrane proteins and proteins involved in biofilm formation and cell motility (19–25). All of the previously validated targets are regulated through binding of the conserved 5’ end region of OmrA/B to their mRNAs. Overall, OmrA and OmrB appear to be important for the restructuration of the bacterial surface in acidic or high osmolarity environments, i.e., in conditions where they are expressed through transcriptional control by the EnvZ-OmpR two-component system (19, 26, 27). Like several known OmrA/B-targets such as *cirA*, f*ecA* and *fepA*, *btuB* encodes a TonB-dependent receptor. However, there are currently no available data showing whether OmrA and OmrB directly or indirectly regulate *btuB* expression and how their cellular function might be coordinated with the regulatory activity of the AdoCbl riboswitch.

**Figure 1.**
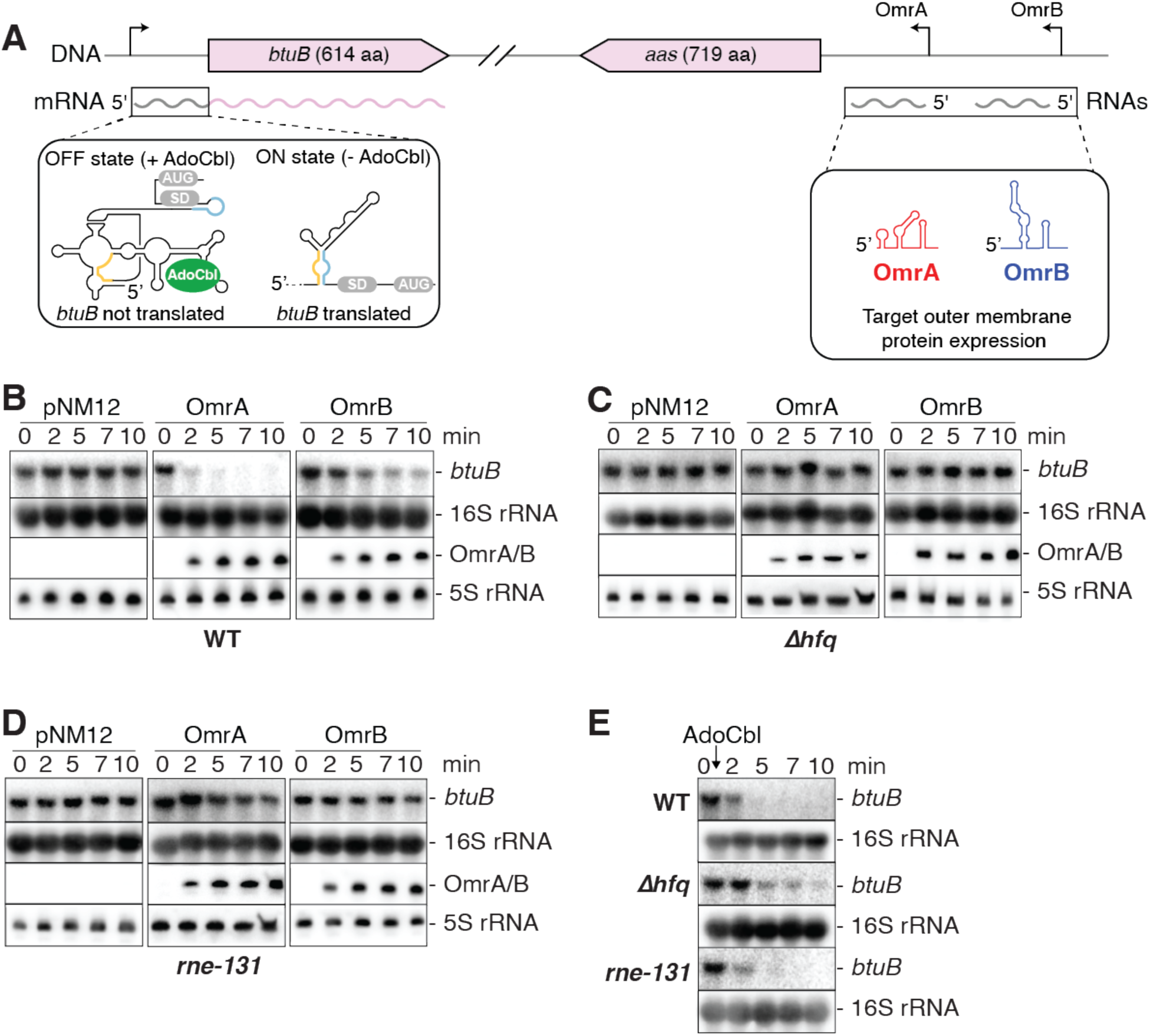
The OmrA and OmrB sRNAs regulate the expression of *btuB.* **(A)** Genomic location of the *btuB* riboswitch and OmrA/B. The left inset shows the ON and OFF riboswitch states when free or bound to adenosylcobalamin (AdoCbl), respectively. OmrA and OmrB are shown in red and blue, respectively. **(B-D)** Northern blot analysis of *btuB* mRNA levels when expressing OmrA or OmrB. The experiments were performed in *E. coli* wild-type (WT) **(B)**, Δ*hfq* **(C)** and *rne-131* **(D)** strains. Total RNA was extracted at the indicated times immediately before (0) or (2, 5, 7 and 10 min) after induction of OmrA or OmrB with 0.1% arabinose. pNM12 is the empty vector control used in these experiments. Specific probes were used to detect *btuB*, OmrA or OmrB. The 16S and 5S rRNA were used as loading controls when monitoring the expression of *btuB* or OmrA/B, respectively. **(E)** Northern blot analysis of *btuB* mRNA levels in the WT, Δ*hfq* and *rne-131 E. coli* strains. Total RNA was obtained at the indicated time relative to the addition of AdoCbl. The 16S rRNA was used as a loading control.

Here, we characterize the genetic regulation exerted by the AdoCbl riboswitch and OmrA/OmrB on *btuB* expression. Our data indicate that *btuB* is repressed by OmrA—and to a lesser extent by OmrB—at the level of translation by promoting binding of the Hfq chaperone to the *btuB* translation initiation region. This sRNA control of *btuB* is not only independent of the riboswitch regulation but also occurs via a distinct mechanism, even though both regulators primarily target *btuB* translation initiation. To our knowledge, our study provides the first example of a mechanism where the regulation of gene expression is achieved through both riboswitch and sRNA control.

## Results

### OmrA/B sRNAs and an AdoCbl-dependent riboswitch negatively control *btuB* mRNA levels

To study the molecular basis of OmrA/B effects on *btuB* expression, we first analyzed by Northern blot assays the levels of endogenous *btuB* mRNA in a strain expressing OmrA or OmrB from a plasmid using an arabinose-inducible promoter (P_BAD_). We performed these experiments in WT, Δ*hfq* and *rne-131* mutant strains, the last of which contains an RNase E variant lacking the C-terminal domain. This mutation prevents the assembly of the RNA degradosome complex (28) and was found to strongly limit the regulation of several mRNAs targeted by OmrA/B (20). In the WT strain, induction of OmrA resulted in a large decrease in *btuB* mRNA levels (Figure 1B), consistent with previous microarray data (19). Induction of OmrB also led to a smaller, but clearly visible reduction in *btuB* mRNA (Figure 1B). In contrast, *btuB* mRNA levels remained largely unaffected when OmrA/B were induced in Δ*hfq* (Figure 1C) and *rne-131* mutant strains (Figure 1D), showing that both Hfq and the RNA degradosome are important for OmrA/B regulation of *btuB* mRNA levels. Of note, the levels of the OmrA/B sRNAs are strongly decreased in an *hfq* null strain ((20, 22), and see later), which may explain the loss of regulation in the Δ*hfq* background. Together, these results show that OmrA and, to a lesser extent OmrB, reduce *btuB* mRNA levels and rely on Hfq and the RNA degradosome to achieve genetic regulation at the mRNA level.

Previous studies have shown that AdoCbl sensing by the *btuB* riboswitch modulates both translation initiation (6, 13, 16, 29) and mRNA levels (13, 30). In agreement with this(12, 26), the addition of AdoCbl promoted a decrease in *btuB* mRNA levels in the WT strain (Figure 1E). In contrast to OmrA/B regulation, we found that this AdoCbl-dependent decrease in mRNA levels was not affected in either Δ*hfq* or *rne-131* mutant strains (Figure 1E), showing that Hfq and the degradosome are not involved in the riboswitch regulation. These data indicate that the AdoCbl riboswitch relies on a different molecular mechanism than OmrA/B to regulate the levels of the *btuB* mRNA. Using a *btuB-mScarlet* fusion as a readout for *btuB* expression (described later with Figure 4), we also found that deleting the gene for the 3’-5’ exoribonuclease PNPase caused a decrease in the OmrA-mediated control of *btuB* as repression dropped from 2.5-fold in the WT strain to 1.4-fold in the *pnp* mutant (Supplementary Figure S2, panels A-C). This is consistent with the known requirement of PNPase for several other sRNA-dependent regulations (31, 32). In contrast, regulation by 1 µM or 5 µM AdoCbl was about 5-fold in the WT and more than 8-fold in the *pnp* mutant (Supplementary Figure S2, panels D-F), thus confirming that sRNA and riboswitch control of *btuB* are most likely due to different mechanisms.

### sRNAs and riboswitch regulation requires different regions of the *btuB* mRNA

To further characterize the regulation of *btuB* by OmrA/B, we next employed translational BtuB-LacZ fusions (Figure 2A) containing the *btuB* 5’ UTR and increasing portions of *btuB* coding sequence (CDS) fused in frame to the 10th codon of *lacZ*. These fusions were expressed under the control of the P_BAD_ arabinose-inducible promoter to ensure that the observed effects did not originate from an endogenous promoter control. In these experiments, OmrA/B were overexpressed from a plasmid with an IPTG-inducible promoter. Our data revealed that OmrA represses by at least 50% the expression of translational fusions containing 81 nucleotides (nt) or more of the CDS (Figure 2B). Stronger OmrA effects were observed when longer regions of the *btuB* CDS were used: the expression of BtuB_120_-, BtuB_210_- and BtuB_420_-LacZ fusions were reduced by ∼6-, ∼5- and ∼5-fold, respectively (Figure 2B), indicating that full regulation requires more than the first 81 nt of *btuB* CDS. Overproduction of OmrB showed a similar trend, but with a systematically lower efficiency, as the maximal repression reached only 50% for the BtuB_120_-, BtuB_210_- and BtuB_420_-LacZ translational fusions (Figure 2C). With a similar strategy, we next investigated the region of *btuB* necessary to regulate mRNA levels by using transcriptional *btuB-lacZ* fusions (Figure 2A). These constructs were designed with the same regions of *btuB* mRNA, followed by a stretch of 4 codons ending with a UAA stop and fused to *lacZ* containing its own translation initiation signals. We observed that only the longest fusion, carrying 420 nt of the CDS, was marginally affected by OmrA and OmrB (reduction of *lacZ* expression by only ∼33% and ∼20%, respectively), while none of the shorter fusions were repressed by the sRNAs (Figures 2B and 2C). Together, these data confirm that OmrA represses *btuB* expression more efficiently than OmrB, consistent with their effect on *btuB* mRNA levels (Figure 1C). Furthermore, a smaller region of *btuB* CDS was required to control the expression of the translational fusions when compared to the transcriptional fusions. These results strongly suggest that these sRNAs primarily act at the translational level and that the decrease in *btuB* mRNA likely results from RNase E cleavage of the messenger devoid of ribosomes, downstream of the 210th nt of the *btuB* CDS.

**Figure 2.**
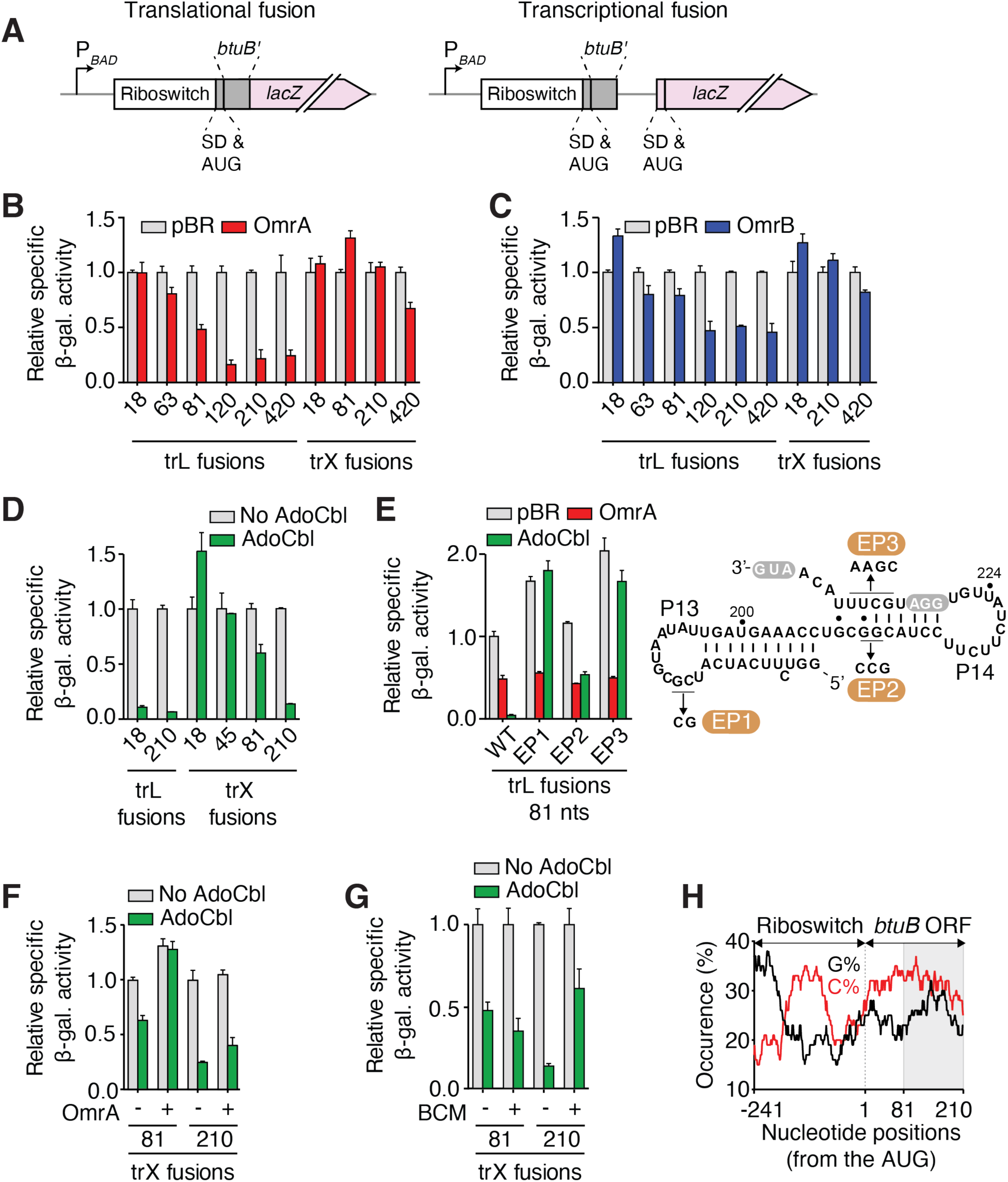
sRNAs and riboswitch regulation rely on different regions of the *btuB* mRNA. **(A)** Schematics showing translational (left) and transcriptional (right) fusions containing various lengths of *btuB* coding region (*btuB’*). An arabinose inducible promoter (P_BAD_) is used to express constructs. The Shine-Dalgarno (SD) and AUG are indicated. **(B-D)** Beta-galactosidase assays of translational BtuB-LacZ and transcriptional *btuB-lacZ* fusions in the presence of OmrA **(B)**, OmrB **(C)** or AdoCbl **(D)**. The number of nucleotides of the *btuB* CDS is indicated for each construct. Values were normalized to the activity obtained in the absence of sRNA (pBR) or without AdoCbl. The average values and the standard deviations were obtained from three independent experiments. **(E)** Beta-galactosidase assays of selected *btuB* mutants destabilizing the stems P13 and P14. Experiments were performed using the BtuB_81_-LacZ translational fusion in the presence of OmrA or AdoCbl. The predicted secondary structure of the region encompassing stems P13 and P14 is shown to the right, and the EP1, EP2 and EP3 mutants are indicated. The SD sequence (GGA) and the AUG are in gray. **(F,G)** Beta-galactosidase assays performed in the absence and presence of AdoCbl when using the *btuB_81_-* and *btuB_210_-lacZ* transcriptional fusions. Experiments were performed either when expressing OmrA **(F)** or when adding bicyclomycin (BCM) **(G)**. **(H)** Sequence analysis of cytosine (%C) and guanine (%G) distribution in the *btuB* sequence. A scanning window of 25 nt was used to determine the C and G occurrences as a function of positions from the AUG start site. The gray region highlights *btuB* region between residues 81 and 210.

We also determined the minimal *btuB* region required for riboswitch regulation. Our reporter gene assays revealed that only 18 and 210 nt of *btuB* CDS are sufficient to trigger strong AdoCbl-dependent regulation of translational and transcriptional constructs, respectively (Figure 2D), in complete agreement with a previous study (13). These results indicate that riboswitch control requires a much smaller region of *btuB* CDS than sRNA control and confirm that regulation by the riboswitch does not depend on the sRNAs. We further investigated the importance of several 5’-UTR secondary structure elements for riboswitch and OmrA-mediated regulation. Since both regulators modulate the expression of the translational BtuB_81_-LacZ fusion (Figures 2B and 2D), we used this construct to characterize the role of two structural elements (helices P13 and P14) that are involved in riboswitch-dependent translation regulation (13, 15, 16, 33). While helix P13 is part of a pseudoknot structure located in the riboswitch aptamer domain, helix P14 directly modulates the access of ribosomes to the SD (Figure 2E) (34). As expected, AdoCbl strongly repressed the WT fusion (by 97%, Figure 2E), and this regulation was lost when the pseudoknot was destabilized via the EP1 mutant. In contrast, repression by OmrA was not affected in the EP1 mutant since it still decreased *btuB* expression by ∼50%, as observed for the WT (Figure 2E). Similarly, the destabilization of the P14 stem (EP2 and EP3 mutants) resulted in loss or decrease of AdoCbl-dependent *btuB* gene repression while retaining OmrA regulation (Figure 2E). Together, these results clearly show that OmrA regulatory activity does not rely on a functional riboswitch to inhibit *btuB* expression.

### OmrA and AdoCbl effectors rely on different regulatory mechanisms

We next assessed whether OmrA binding to *btuB* mRNA could interfere with the AdoCbl riboswitch regulation. To do so, we employed the 81 nt *btuB-lacZ* transcriptional fusion whose expression is not regulated by OmrA, but is efficiently repressed by AdoCbl (Figures 2B and 2D). Remarkably, when OmrA was overexpressed, the effect of AdoCbl was completely abolished (Figure 2F), suggesting that OmrA binding prevents the riboswitch from modulating the mRNA levels of this 81 nt *btuB-lacZ* transcriptional fusion. However, when repeating these experiments using a longer *btuB* transcriptional fusion of 210 nt, the addition of AdoCbl decreased *btuB* mRNA levels even in presence of OmrA (Figure 2F). These results indicate that *btuB* sequence elements located between positions 81 and 210 of the CDS enable a more robust mRNA regulation by the AdoCbl riboswitch, that is no longer affected by OmrA. Therefore, we investigated which regulatory mechanisms are at play in this region. We reasoned that, in addition to the known translational repression, the regulation could involve the transcription termination factor Rho, as previously determined for several riboswitches (6). Accordingly, we monitored the regulation of *btuB* expression in the presence of the Rho inhibitor bicyclomycin (BCM)(35). We found that, while BCM does not perturb AdoCbl regulation of the 81 nt construct, it relieved AdoCbl-dependent repression by ∼4-fold in the 210 nt fusion (Figure 2G). Hence, following AdoCbl-dependent translation inhibition, Rho could target the untranslated 81-210 nt region of the *btuB* mRNA to terminate transcription. In agreement with this hypothesis, this region of *btuB* is characterized by a C-rich and G-poor sequence (Figure 2H) that is often associated with the presence of Rho utilization (*rut*) sites (36, 37). Together, our results suggest that the AdoCbl riboswitch and OmrA modulate *btuB* mRNA levels using different mechanisms that exploit specific *btuB* sequences to allow Rho-dependent transcription termination (for riboswitch control), and RNA decay mediated by RNase E and the degradosome (for sRNA control).

### OmrA recognizes *btuB* mRNA via its specific central region

To decipher how OmrA could control *btuB* expression, we used the IntaRNA *in silico* prediction tool (38) to search for a potential base-pairing interaction between OmrA and the first 100 nt of the *btuB* CDS, and extended the prediction by hand (Figure 3A). The *btuB*-OmrA interaction is predicted to rely mostly on the central region of OmrA, which diverges from the OmrB sequence, in complete agreement with the different regulation efficiencies observed for the two sRNAs (Figures 1B, 2B and 2C). To validate this prediction, different mutations were introduced in OmrA and in a translational fusion of *btuB* mRNA region (−240 to +99) followed by *lacZ* coding sequence only a few nts downstream of the pairing prediction (BtuB_99_-LacZ). This fusion is transcribed from a constitutively expressed P_LtetO-1_ promoter. We first found that mutating positions 26-36 of OmrA (OmrAM9* mutant) negatively affected the regulation, since the WT fusion was more efficiently repressed by OmrA (5-fold) than by OmrAM9* (1.6-fold) (Figure 3B). Similarly, mutating *btuB* positions +71 to +78 (*btuB*M9 mutant) strongly impaired control by the WT OmrA sequence (1.2-fold repression), in agreement with this sequence being important for OmrA recognition. However, combining both the OmrAM9* and *btuB*M9 mutants to restore the predicted OmrA-*btuB* interaction only slightly improved repression (1.7-fold) compared to the mutant fusion/WT OmrA pair, and did not lead to significant gain in regulation compared to the WT fusion/OmrAM9* pair (Figure 3B). This could be explained by the presence of the unpaired residues 32-34 and 38-41 in OmrA (Figure 3A), adjacent to the mutated positions in the M9* variant and possibly unfavourable for compensation. Consistent with this, removing these unpaired nucleotides in the OmrAM9* context restored control of the *btuB*M9 variant (see OmrAopt and OmrAoptM9* mutants in Supplementary Figure S3).

**Figure 3.**
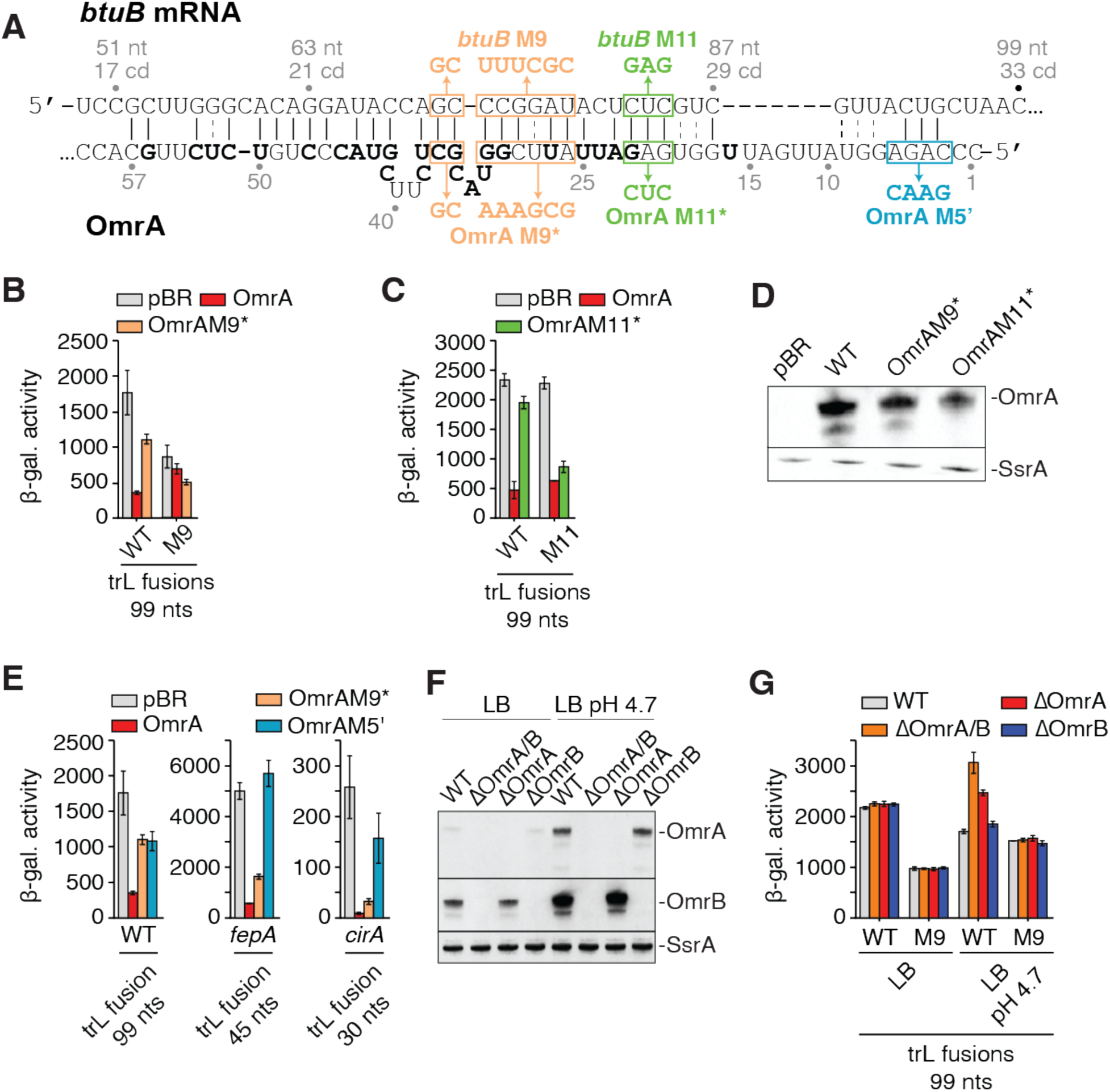
OmrA regulates *btuB* expression through direct interaction via its central region. **(A)** Predicted base-pairing between *btuB* and OmrA. The numbering scheme is relative to the *btuB* start codon and the OmrA transcription start site. The non-conserved nucleotides between OmrA and OmrB are shown in bold. The regions containing the mutations to investigate the OmrA-*btuB* interaction are shown in boxes. **(B, C, and E)** Beta-galactosidase assays of BtuB_99_-LacZ translational fusions carrying the M9 mutations **(B)** or the M11 mutations **(C)**, and of FepA_45_-LacZ and CirA_30_-LacZ translational fusions **(E)** upon overproduction of different OmrA variants. All fusions are expressed from a P_LtetO-1_ constitutively expressed promoter. Strains used in these assays are deleted of *omrA* and *omrB* chromosomal copies. The β-galactosidase average values and the standard deviations were obtained from three independent experiments. **(D)** Northern blot analysis of levels of OmrA M9* and M11* variants using RNA extracted from cell cultures used for corresponding ß-galactosidase assays. Detection of the SsrA RNA was used as a loading control. **(F)** Northern blot analysis of OmrA and OmrB levels in LB and in LB pH 4.7. Experiments were done in cells carrying or not the chromosomal *omrA* and/or *omrB* genes as indicated and RNA was extracted from the same cultures that were used for beta-galactosidase assays of WT BtuB_99_-LacZ in **(G)**. The SsrA RNA was used as a loading control. **(G)** Beta-galactosidase assays of BtuB_99_- LacZ and BtuB_99_M9-LacZ translational fusions in cells carrying or not the chromosomal *omrA* and/or *omrB* genes, in LB and in acid LB (pH 4.7). Shown are the average β-galactosidase activities and standard deviations of three independent experiments.

**Figure 4.**
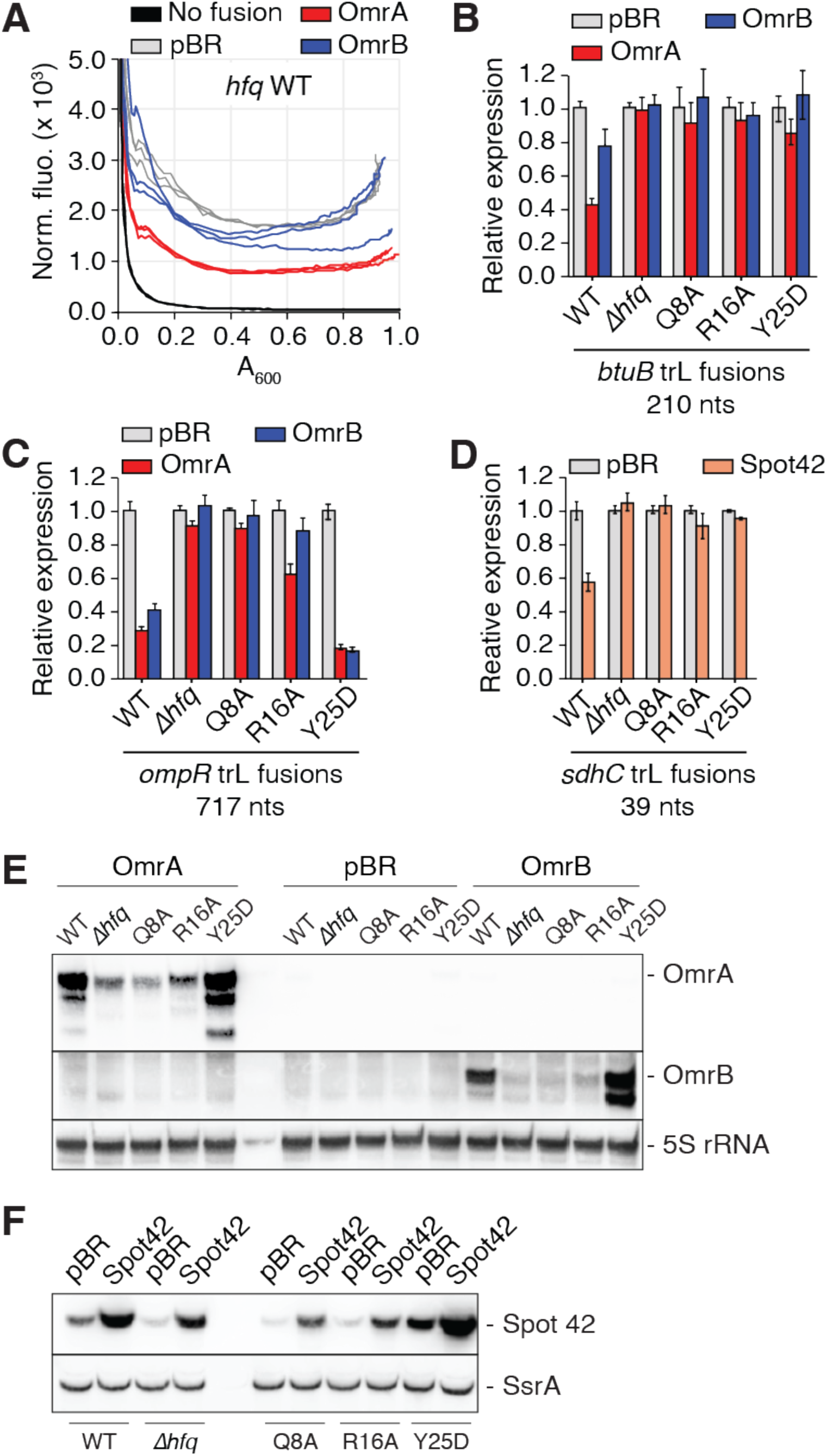
The Hfq distal face is required for the control of *btuB* by OmrA. **(A)** Representative fluorescence measurement of a BtuB_210_-mScarlet translational fusion upon overproduction of OmrA or OmrB over a 16 h time window. Shown is the normalized fluorescence that corresponds to the fluorescence divided by the absorbance at 600 nm, as a function of the absorbance at 600 nm. A control was performed using an isogenic strain lacking the fluorescent construct and carrying the pBRplac (pBR) empty plasmid (’no fusion’). **(B-D)** Fluorescence assays of BtuB_210_- **(B)**, OmpR_717_*-* **(C)** and SdhC_39_-mScarlet **(D)** translational fusions upon overproduction of OmrA or OmrB. Experiments were performed in isogenic *hfq+* (WT), Δ*hfq*, *hfqQ8A*, *hfqR16A* and *hfqY25D* strains. Measurements were done in triplicate and shown is the fluorescence normalized to the absorbance at 600 nm, at an absorbance at 600 nm close to 0.3 and set arbitrarily at 1.0 for the WT strain transformed with the vector control. **(E,F)**. Northern blot analysis of levels of OmrA and OmrB **(E)** and Spot42 **(F)** in isogenic *hfq+* (WT), Δ*hfq*, *hfqQ8A*, *hfqR16A* and *hfqY25D* strains. The 5S rRNA and the SsrA RNA were used as loading controls. Total RNA was extracted from the same set of strains that were used in panels **B** and **D**, respectively. A longer acquisition of the OmrA and OmrB Northern-blots of panel **E** is provided in Supplementary Figure S9.

We thus used an additional pair of mutants to assess the OmrA-*btuB* interaction (mutants *btuB*M11 and OmrAM11*, Figure 3A). The mutation *btuB*M11 is not sufficient to prevent OmrA regulation, which is most likely explained by the existence of alternative pairing in this case. Nonetheless, the OmrAM11* variant no longer repressed the WT BtuB-LacZ fusion, but was efficient in regulating the BtuBM11-LacZ fusion (∼2.6-fold repression, Figure 3C), despite the lower levels of OmrAM11* compared to OmrA WT (Figure 3D). Thus, taken together, the M9 and M11 sets of mutants strongly support the hypothesis that the central region of OmrA interacts with residues +66 to +79 of the *btuB* CDS *in vivo*.

This base-pairing is highly unusual compared to previously reported targets of the OmrA/B sRNAs such as *cirA*, *fepA*, *ompR*, *ompT*, *csgD, flhDC*, *dgcM* and *flgM* mRNAs that all interact with the conserved 5’ end of OmrA/B (20–22, 24, 25, 39). This specificity of the OmrA-*btuB* interaction was further examined by comparing the effects of mutations in OmrA 5’ end or central region (Figure 3A, see M5’ and M9* mutants, respectively) on *btuB* or *cirA* and *fepA*, two known targets of OmrA 5’-end. The OmrA M5’ mutation alleviated the repression of BtuB_99_-LacZ fusion (Figure 3E) similar to what we observed with OmrA M9*. Hence, neither mutation alone abolished regulation completely, which could be explained by the length of the predicted interaction between OmrA and *btuB*, that may also involve pairing of OmrA 5’-end with *btuB* residues +83 to +94 (Figure 3A). In contrast, translational fusions FepA_45_-LacZ and CirA_30_-LacZ, were efficiently repressed by OmrAM9* but very poorly regulated by OmrAM5’ (Figure 3E), as expected with classical OmrA 5’-end targets. These data confirm the crucial and unique role played by the OmrA-specific central region in the control of *btuB*.

### Physiological levels of OmrA and OmrB regulate *btuB* expression

We next analyzed the role of OmrA/B in *btuB* regulation under more physiological conditions, i.e., when the sRNAs are expressed from their native chromosomal locus. For these assays, we followed the activity of the BtuB_99_-LacZ translational fusion in an *omrAB+* strain, or in strains deleted for *omrA*, *omrB* or both. These experiments were performed in standard LB medium (measured pH of 6.8), or in acidified LB (pH 4.7) where OmrA/B expression is expected to be strongly induced through OmpR activation (27, 40), which was confirmed by Northern blot (Figure 3F). No difference in expression of the WT BtuB_99_-LacZ fusion was observed in the presence or absence of OmrA and/or OmrB in standard LB (Figure 3G), consistent with the relatively low levels of OmrA/B in this condition (Figure 3F) (27, 40). In contrast, when these experiments were performed in acidified LB, the activity of the WT fusion was increased by ∼1.9-fold and ∼1.5-fold in the Δ*omrAB* and Δ*omrA* deletion strains, respectively (Figure 3G), confirming the repression of *btuB* expression by physiological levels of OmrA. No such effect was observed with the sole deletion of OmrB (Figure 3G), further supporting that *btuB* is a preferential target of OmrA over OmrB. Finally, the expression of the M9 mutant version of the BtuB_99_-LacZ fusion was not significantly affected in any of the *omrA/B* deletion strains (Figure 3G), in agreement with the direct base-pairing interaction shown in Figure 3A.

### The distal face of Hfq is required for *btuB* regulation by OmrA

Given that the OmrA pairing site is within the *btuB* CDS, downstream of the translation initiation region (TIR), we next investigated the molecular mechanism by which OmrA represses *btuB* expression. We first analyzed the role of Hfq in this sRNA regulation by taking advantage of previously characterized point mutants in three different RNA-binding surfaces of Hfq: the proximal face (mutant Q8A), the rim (R16A) and the distal face (Y25D) (41, 42). The proximal face and the rim are important for binding many Hfq-dependent sRNAs, also called Class I sRNAs, while the distal face has been implicated in binding successive A-R-N (R=purine, N=any nucleotide) motifs present in the mRNA targets of such sRNAs (41). Of note, OmrA and OmrB display the features of Class I sRNAs as their levels are strongly decreased in the proximal and rim mutants, but not in the distal face mutant (24).

To assess the role of Hfq in the regulation of *btuB* expression, we employed a BtuB_210_- mScarlet-I (*mSc*) translational fusion and monitored mSc fluorescence upon overexpression of OmrA in strains carrying one of the following five *hfq* alleles: WT hfq, Δ*hfq*, Q8A, R16A, or Y25D. As a control, we also monitored the repression of *ompR* and *sdhC*, which are controlled by OmrA/B (20) and Spot42 (43), respectively, using translational mSc fusions in the same *hfq* backgrounds. Importantly, *ompR* is a canonical OmrA/B target because the sRNA pairing site overlaps with the TIR, most likely allowing direct competition with the binding of the ribosomal 30S subunit (20). In contrast, *sdhC* is repressed by Spot42 sRNA in a non-canonical manner: the sRNA pairs upstream of the TIR and recruits Hfq to the TIR, thus allowing translation repression (43).

In the *hfq* WT background, the fluorescence of the BtuB-mSc fusion was decreased more than 2-fold upon OmrA overproduction, while OmrB had a marginal effect (∼1.3-fold) (Figures 4A and 4B and Supplementary Figure S4). In the same conditions, the expression of the *ompR-mSc* fusion was decreased more than 3- and 2-fold, respectively (Figure 4C and Supplementary Figure S5). The sRNA-dependent regulation of both *btuB* and *ompR* was abolished, or strongly reduced, in the absence of Hfq (Δ*hfq*) or when mutated in the proximal (Q8A) or rim (R16A) faces (Figures 4B and 4C). This is explained, at least in part, by the much lower accumulation of OmrA/B in these mutants (Figure 4E). In contrast, OmrA and OmrB accumulated to higher levels in the context of the Hfq distal face mutant (Y25D) (Figure 4E), as observed previously (24). In this context, the expression of the *btuB-mSc* fusion was no longer regulated by OmrA or OmrB (Figure 4B), while *ompR-mSc* was still repressed more than 5-fold by both sRNAs (Figure 4C). The regulation of *ompR* in the Y25D mutant strain indicates that OmrA/B do not rely on the Hfq distal face to regulate *ompR*, consistent with the model that these sRNAs bind directly to the *ompR* TIR and compete with the 30S ribosomal subunit (26). In contrast, the lack of regulation of *btuB* in the *hfq* Y25D mutant suggests that OmrA/B may instead use a different mechanism in this case, in which Hfq binding to the mRNA is required. Such a non-canonical control mechanism was observed for instance in the modulation of *sdhC* expression by Spot42 (43) or of *manX* by SgrS and DicF sRNAs (44), with the sRNAs recruiting Hfq to the TIR for translation inhibition. Interestingly, the repression of *sdhC* and *btuB* by their respective sRNAs was similarly strongly reduced in all four *hfq* mutants (Figures 4B and 4D and Supplementary Figure S6), hinting that the regulatory mechanism involved in both cases could be similar. In support of an important role of Hfq in *btuB* control, we also found that *btuB* expression was increased in the Δ*hfq* or in the *hfq*Y25D strain (Supplementary Figure S7), while *hfq* overexpression from a plasmid (45) led to *btuB* repression (Supplementary Figure S8). Again, the same pattern was observed with the expression of *sdhC*, while no strong effect of Hfq was visible on *ompR* in these experiments.

### Hfq interacts with OmrA and the *btuB* translation initiation region *in vitro*

These results prompted us to investigate the interaction between Hfq and the *btuB* mRNA, and its function in *btuB* regulation, in greater detail using *in vitro* approaches. We first performed lead acetate footprinting experiments to identify the Hfq binding site on the *btuB* mRNA, using a *btuB* mRNA fragment ranging from nts −80 to +161. An Hfq-dependent protection was observed between positions −8 to +12 of the *btuB* mRNA, thus overlapping with the *btuB* start codon (Figure 5A). Importantly, this protected region contains repetitive A-R-N motifs that are consensus binding sites for the Hfq distal face. We thus tested *in vivo* the effect of a mutation that eliminates these A-R-N motifs on the control by OmrA. The introduction of three CC pairs within the Hfq binding site of the BtuB_99_-LacZ fusion (Figure 5F, H1 mutant) strongly reduced the regulation by OmrA, ∼5.8-fold and ∼1.3-fold for the WT and H1 mutant (Figure 5B), respectively. However, this H1 change also strongly decreased the expression of the BtuB_99_-LacZ fusion. To rule out the possibility that the reduced repression of this H1 mutant by OmrA is due to its lower expression, we verified that a low expression of *btuB*, for instance with a mutation in the P_LtetO-1_ promoter (lowPtet), still allowed OmrA regulation (Figure 5B, ∼4.2-fold). Combined, these data strongly suggest that OmrA regulation of *btuB* requires Hfq and its binding in the vicinity of the TIR to repress *btuB*.

**Figure 5.**
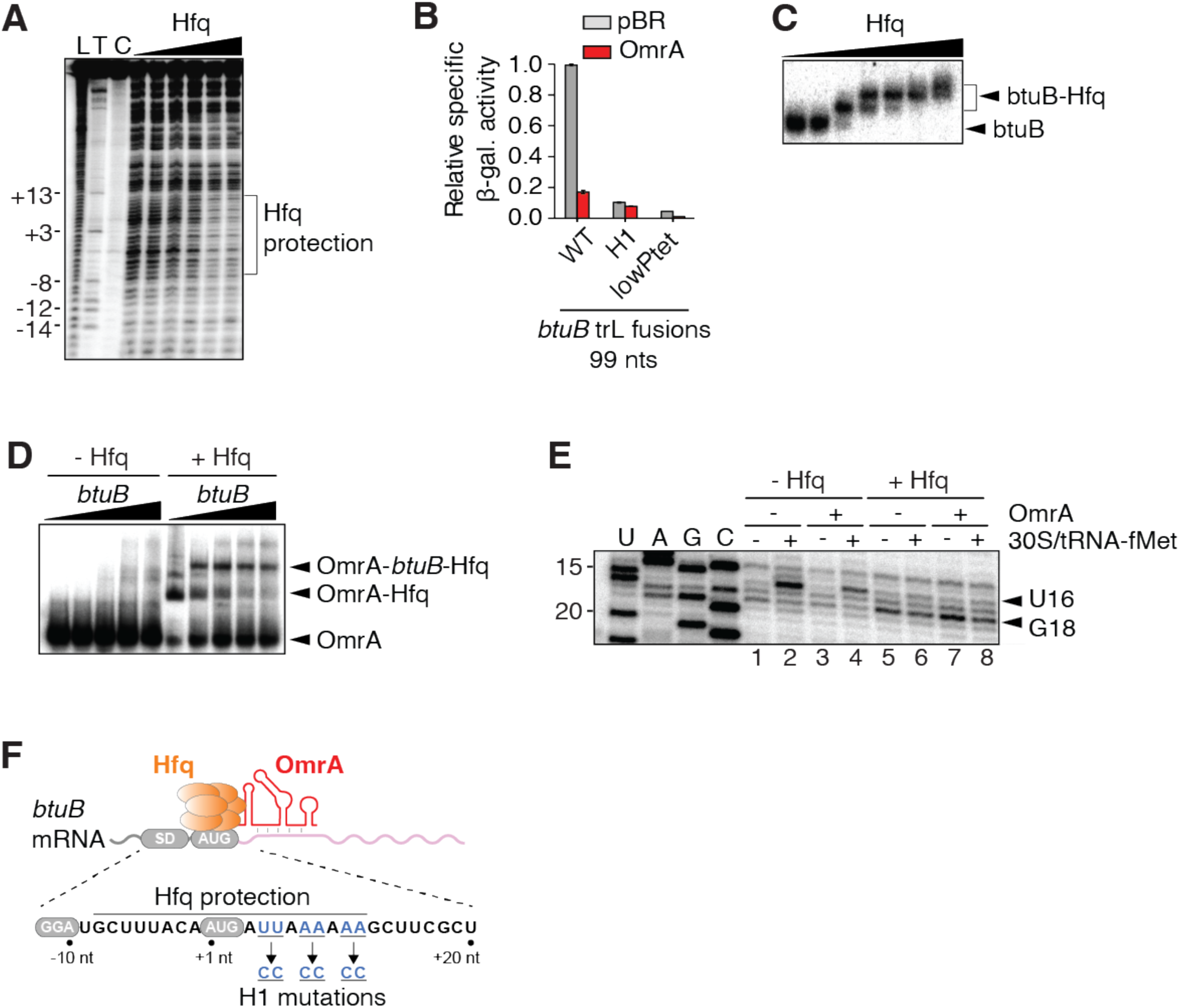
*btuB* mRNA interacts with OmrA and Hfq *in vitro*. **(A)** Lead acetate probing of Hfq binding to the *btuB* mRNA. Radioactively labeled *btuB* RNA corresponding to nts −80 to +161 relative to start codon was incubated with increasing concentrations of Hfq (0 to 20 nM). Lanes L, T and C correspond to an alkaline hydroxide ladder, a partial RNase T1 digestion and a non-reacted control, respectively. The positions relative to the AUG are indicated on the left. **(B)** Beta-galactosidase assays of BtuB_99_-LacZ translational fusions either WT or carrying the H1 mutation in the presence of OmrA. Shown are the average β-galactosidase activities and standard deviations of two independent experiments. **(C)** EMSA assays performed using radiolabeled *btuB* RNA with increasing concentrations of Hfq. Arrows on the right of the gel indicate the free *btuB* RNA and the *btuB*-Hfq complex. The presence of multiple species when increasing Hfq concentration suggests that Hfq may have multiple *btuB* RNA binding sites. **(D)** EMSA assays performed with radiolabeled OmrA incubated with increasing concentrations of *btuB* RNA in the absence or presence of Hfq. The arrows on the right show the free OmrA and the OmrA-Hfq-*btuB* complexes. **(E)** Toeprinting assays monitoring the effect of Hfq and OmrA on the formation of the translation initiation complex. Experiments were performed in the absence or presence of Hfq, OmrA or 30S/tRNA-fMet. Lanes U, A, G and C represent sequencing ladders. **(F)** Scheme representing the *btuB* mRNA with Hfq and OmrA. The Hfq protected region, the Shine-Dalgarno (SD) and AUG are shown. Nucleotides changed in the H1 mutant are shown in blue.

We next used electrophoresis mobility shift assays (EMSA) to characterize the interaction between *btuB*, OmrA and Hfq. Our results showed that a bi-partite *btuB-*Hfq complex is formed upon increasing the concentration of Hfq (Figure 5C). Furthermore, the *btuB-*OmrA interaction was only detected in the presence of Hfq (Figure 5D), indicating that Hfq is required for the formation of a stable *btuB*-OmrA complex. These results suggest a regulatory mechanism in which OmrA and Hfq interact directly with *btuB* to modulate its genetic expression.

To directly assess the role of Hfq in inhibiting translation initiation, we used *in vitro* toeprint assays in the presence or absence of Hfq (Figure 5E). The assay was performed using the 30S ribosomal subunit, *btuB* mRNA and the initiator tRNA (tRNA-fMet) (46). In the presence of 30S and tRNA-fMet, a reverse transcription stop (toeprint) was detected at position U16 (Figure 5E, lane 2), indicating that the 30S subunit is bound to the start codon of the *btuB* mRNA, in agreement with previous data (34). In the presence of Hfq, a clear loss of the U16 toeprint was observed (Figure 5E, lane 6), showing that Hfq prevents the 30S subunit from binding the *btuB* mRNA. Instead, a new reverse transcription stop was observed at position G18, likely due to Hfq blocking the reverse transcriptase at this position, in good agreement with our lead probing data (Figure 5A). When the experiment was performed with OmrA, the addition of Hfq still resulted in the loss of U16 toeprint and the generation of the G18 signal (Figure 5E, lanes 4 and 8), corresponding to Hfq binding.

Together, these results demonstrate that Hfq binds directly to *btuB* mRNA upstream of the OmrA pairing site and prevents the 30S subunit from binding to the *btuB* AUG start codon *in vitro* (Figure 5E). This, coupled with the requirement for Hfq for OmrA-mediated regulation *in vivo* (Figures 1C, 4B and 5B), suggests that binding of OmrA helps recruit Hfq to the *btuB* TIR, similar to the way Spot42 or SgrS/DicF recruit Hfq to regulate *sdhC* or *manX* expression, respectively (43, 44).

## Discussion

The findings reported here identify *btuB* as a new member of the Omr regulon. Even though the physiological role of these two sRNAs is still not fully understood, *btuB* is reminiscent of other previously validated targets of OmrA/B. First, BtuB is an outer membrane protein (OMP), like several other known targets (OmpT, CirA, FecA and FepA). Second, like CirA, FecA and FepA, BtuB depends on the TonB system for the uptake of its substrate, namely AdoCbl.

Importantly, there is one notable difference between *btuB* and the other OmrA/B targets. While the pairing of OmrA/B to all previously recognized targets relies on the sRNAs 5’ end (20–22, 24, 25, 39) and our unpublished data for *fecA*), control of *btuB* involves the central region of OmrA, that distinguishes it from OmrB. Consistent with this, deletion of OmrA is sufficient to increase *btuB* expression in acid medium, while under the same conditions, the deletion of OmrB alone had no effect (Figure 3G). However, in the absence of OmrA, deletion of OmrB allowed a further increase in *btuB* expression, but not that of *btuB* carrying the M9 mutation (Figure 3G), suggesting that OmrB can still pair to *btuB* mRNA, although with a much lower efficiency than OmrA. In this regard, a 9 bp complementarity exists between residues 29 to 37 of OmrB and +69 to +77 of *btuB*, but whether this participates in regulation is still unclear. Furthermore, it is possible that, similarly to OmrA (Figure 3A), OmrB targets the +85 to +94 *btuB* region through its 5’ domain. However, more work would be required to ascertain this regulatory interaction. In sum, *btuB* appears as a preferential OmrA target, while modulation by OmrB is only marginal and possibly requires more specific conditions.

To our knowledge, this is the first report of a preferential OmrA target via pairing to the OmrA specific region, indicating that, in addition to a common regulon, some genes are likely to be regulated by OmrA alone. The reverse could also be true, *i.e.*, that some genes could be controlled by OmrB alone, even though no such examples have been described so far. Differential control of targets such as reported here for *btuB* is likely to have contributed to the conservation of both OmrA and OmrB sRNAs in enterobacteria.

The stimulation of OmpR activity in acidic conditions (27, 40) provides a rationale to explain the induction of OmrA/B following acid stress (Figure 3G). In such low pH conditions, it has been reported that the proton motive force is increased (47, 48), the latter being crucial for cobalamin import by BtuB (49) and the function of other TonB-dependent transporters. These results are consistent with a model in which acidic conditions promote the import of cobalamins through BtuB, ultimately leading to *btuB* repression through both AdoCbl binding to the riboswitch and OmrA/B activity. Furthermore, *btuB* transcription is inhibited at the promoter level by the transcriptional regulator GadX, whose synthesis is induced under acid stress (50). Globally, with the above results, our data show that *btuB* expression is highly regulated at multiple levels by GadX, the AdoCbl riboswitch and the sRNAs OmrA/B.

From a mechanistic perspective, Hfq is strictly required for the control of *btuB* by OmrA/B. This was expected and has been observed for many of the other Omr targets since OmrA/B levels significantly drop in the absence of this chaperone (Figure 4E)(20, 22, 24). More unusual, however, is the fact that this control was also abolished in an Hfq distal face mutant (Y25D) as well. This is not generally observed for canonical regulation where sRNAs pair to the RBS (such as *ompR* control by OmrA/B, Figure 4C). Because the Hfq distal face has been shown to be involved in mRNA binding, this strongly indicates that OmrA control of *btuB* is dependent on the ability of Hfq to interact with this target mRNA. Interestingly, the same is true for the control of *sdhC* by Spot42 (Figure 4D) and of *dgcM* by OmrA/B (24). In the latter case, Hfq was found to mediate a change in the structure of *dgcM* mRNA facilitating the interaction with OmrA/B (24), whereas in the former case, as mentioned previously, Hfq is directly responsible for the inhibition of 30S binding to *sdhC* (43). Our *in vitro* experiments showed that Hfq binds to the TIR of *btuB* mRNA, in a region containing a canonical binding site formed by multiple (A-R-N) motifs, thereby inhibiting 30S binding (Figure 5E). In contrast, OmrA by itself had a much weaker effect on toeprint formation (Figure 5E). Together, these results suggest a mechanism reminiscent of that described for the Spot42-*sdhC*, SgrS-*manX* or DicF-*manX* pairs, where the central region of OmrA binds with *btuB* mRNA, thereby promoting Hfq recruitment and inhibiting 30S binding. Furthermore, Hfq significantly improves the formation of an OmrA-*btuB* complex *in vitro* (Figure 5D). The precise mechanism underlying this observation is still unclear at this stage, but Hfq could be involved in the remodeling of OmrA or *btuB* mRNA structure to facilitate formation of the duplex, similarly to what has been reported for *dgcM* (24).

While most studies performed in the last decades have led to the idea that sRNAs regulate gene expression at the post-transcriptional level (51–53), it has recently been shown that sRNAs can also operate cotranscriptionally by controlling Rho transcription termination (54–56). For ChiX, Rho transcription termination is indirectly controlled through the modulation of translation initiation in *Salmonella enterica* (57). However, for several sRNAs such as DsrA, ArcZ and RprA, Rho activity is directly modulated by sRNA association to the mRNA, presumably by inhibiting Rho binding or translocation (7). In contrast, current data indicate that riboswitches mostly regulate gene expression at the co-transcriptional level both in *Bacillus subtilis* (58, 59) by modulating intrinsic transcription terminators, and in *E. coli* by relying on Rho to downregulate mRNA levels following translation inhibition (6). It was also shown that post-transcriptional regulation may be used by some riboswitches (59), suggesting that like sRNAs, riboswitches may exert control both during and after the completion of the transcriptional process.

It is remarkable that translation initiation of *btuB* is selectively modulated by both the AdoCbl riboswitch and OmrA/B sRNAs, which affect mRNA levels through different mechanisms. As the riboswitch-mediated decrease in *btuB* mRNA is independent of both Hfq and the degradosome (Figure 1), this strongly suggests that blocking translation initiation of *btuB* is not sufficient to lead to RNase E-mediated decay of this mRNA. Instead, the degradation that is observed upon OmrA/B overexpression most likely relies on the recruitment of RNase E and the degradosome to the *btuB* mRNA through the Hfq-RNA complex (60, 61). Conversely, the fact that much longer regions of *btuB* mRNA are required for OmrA/B control than for riboswitch control indicates that, while the riboswitch action leads to premature Rho-dependent transcription termination, this may not be the case for the sRNAs. Such a mechanistic difference is compatible with the AdoCbl riboswitch and sRNAs OmrA/B regulating *btuB* gene expression respectively at the co-transcriptional and post-transcriptional levels. Additional experiments will now be required to fully understand to which extent these regulatory activities are confined to co-transcriptional and post-transcriptional mechanisms.

It has been previously reported that riboswitches and sRNAs may act in concert to control gene expression (62–64). Indeed, in *Listeria monocytogenes*, transcription termination caused by the *S-*adenosylmethionine riboswitch generates an sRNA that controls the expression of PrfA, which is involved in virulence gene expression (62). Furthermore, it was shown in *Enterococcus faecalis* and *L. monocytogenes* that the AdoCbl riboswitch controls the formation of an sRNA containing ANTAR RNA elements, the latter being important for the regulation of the *eut* genes involved in ethanolamine utilization (63, 64). While these studies revealed that riboswitches might act as "pre-sRNA" regulatory elements, the results presented in our study rather indicate that riboswitches and sRNAs both modulate the expression of *btuB*.

Riboswitches and sRNAs may participate in the regulation of same mRNA populations, thereby increasing the complexity of gene regulation mechanisms. While riboswitches are in most cases restricted to 5’-UTRs, the combination with sRNA regulation that can target multiple mRNA regions could significantly enhance gene regulation efficiency at both the co-transcriptional and post-transcriptional levels, respectively. The great variety of mechanisms through which riboswitches and sRNAs regulate gene expression could therefore be used and combined in bacteria to ensure cellular homeostasis under multiple conditions. In this regard, it is interesting that other *E. coli* mRNAs known to be regulated by riboswitches were found enriched after immunoprecipitation with the RNA chaperones ProQ or Hfq, and that sRNA-mRNA pairs such as MicA-*moaA* or CyaR-*lysC* were identified in the RNA-RNA interactome studied in the Hfq co-IP fraction (65). This suggests that the dual riboswitch-sRNA control of a single gene reported here is not restricted to *btuB* mRNA, and the study of other systems is likely to be instructive in the future.

## Materials and Methods

### Bacterial strains and plasmids

Bacterial strains and plasmids used in this study are described in Supplementary Table S1. Cells were grown in M63 minimal medium containing 0.2% glucose for experiments where AdoCbl concentration was adjusted (Figures 1 and 2), in CAG medium (minimal A salts (66), 0.5% (w/v) glycerol, 0.25% (w/v) casamino acids, 1 mM MgSO_4_) for fluorescence measurements (Figure 4), or in LB for other experiments. These media were supplemented with antibiotics, AdoCbl, arabinose or IPTG as needed (see Supplementary Information for details).

Gene fusions to *lacZ* or *mScarlet* reporters were constructed by recombineering using recipient strains carrying a mini-lambda and a *cat-sacB* allele upstream of the *lacZ* or *mScarlet* genes. These recipient strains are PM1205 (for construction of P_BAD_-driven *lacZ* fusions), MG1508 (P_LtetO-1_-driven *lacZ* fusions) and OK510 (*mScarlet* fusions). Deletions of *omrA*, *omrB*, or both were made by recombineering of a kanamycin-resistance cassette into the *omr* locus; these mutations, as well as Δ*hfq* and *rne-131* alleles, were moved by P1 transduction as necessary. The different *hfq* point mutants were obtained from D. Schu and N. Majdalani (NIH, Bethesda) and moved by P1 transduction as described in Supplementary Information.

OmrA and OmrB sRNAs were overproduced from either pNM12- or pBRplac-derivative plasmids, where their expression is under the control of an arabinose- or IPTG-inducible promoter, respectively. Mutations were introduced into the pBRplacOmrA plasmid by amplification with mutagenic primers, DpnI digestion, transformation into NEB-5 *lacIq* strains, and sequencing of the resulting plasmids.

The sequences of DNA oligonucleotides used in this study are in Supplementary Table S2.

### ß-Galactosidase Assays

The β-galactosidase activity of transcriptional or translational *btuB-lacZ* fusions expressed from a P_BAD_ promoter (Figure 2) was measured in kinetic assays as previously described (5). Briefly, a bacterial culture was grown overnight in M63 0.2% glycerol minimal medium and was diluted 50-fold into fresh medium, which was then incubated at 37 °C until an OD_600_ of 0.1 was obtained. Arabinose (0.1%) was then added to induce the expression of *lacZ* constructs. AdoCbl (5 µM) and/or IPTG (1 mM) to allow OmrA/B induction from a pBR plasmid was added when indicated. When using bicyclomycin (BCM; 25 µg/mL), assays were performed in 3 mL of culture media. The β-galactosidase activity of other *lacZ* fusions (Figure 3 and Supplementary Figure S3) was measured using a standard Miller assay. Briefly, an overnight culture was diluted 500-fold in fresh LB-Tet-IPTG 100 μM medium and cells were grown to mid-exponential phase. Next, 200 μl aliquots were then mixed with 800 μl of Z buffer, and the activity was measured as previously described (67) after cells were lysed with chloroform and SDS.

### Northern Blots Analysis

Total RNA was extracted from cells grown to midlog phase as described in the previous paragraph using the hot phenol method as in (68). When cells were grown in LB, 650 μl of cell culture were directly mixed with phenol, while cells grown in M63 or CAG were first resuspended in the same volume of sterile water and then mixed with phenol. After one phenol/water and two phenol/chloroform extractions, RNA was precipitated and resuspended in water. A constant amount of total RNA was loaded on 1% agarose and transferred by capillarity onto an Hybond N+ (Amersham) membrane to detect *btuB* mRNA. For detecting OmrA, OmrB and Spot42 sRNAs, total RNA was separated on a 5% polyacrylamide gel and electro-blotted to an Hybond N+ membrane. A radiolabeled probe was used to detect *btuB* as in (5) and biotinylated probes were used to detect OmrA, OmrB or Spot42 as previously described (23).

### mScarlet Fluorescence Assays

The protocol for fluorescence measurement is detailed in the Supplementary data. Briefly, a saturated culture was diluted 500-fold in 200 µl of CAG medium supplemented with tetracycline and IPTG 250 µM in a dark 96-well plate with clear bottom. Wells were then covered with 50 µl mineral oil and bacterial growth and fluorescence were followed every 12 minutes during a 16-hours kinetic where the plate was incubated at 37°C with shaking at 500 rpm in a Clariostar plate reader. Absorbance was followed at 600nm, and fluorescence was measured using an excitation wavelength of 560nm and emission at 600nm (with a 15 nm bandwidth). Measurements were systematically done in triplicate and normalized to the absorbance at 600nm. For Northern blot analysis of the sRNA levels in the different *hfq* mutants (Figures 4E and 4F), total RNA was extracted from the same strains and same growth media as those used for fluorescence, but grown to mid-exponential phase in Erlenmeyer flasks rather than in 96-wells plates.

### *In vitro* transcription

*In vitro* transcription reactions were performed as previously described (5). Briefly, PCR products were used as DNA templates and contained a T7 RNA polymerase promoter. After transcription reactions were incubated for 3 hours at 37 °C, RNA products were precipitated with ethanol and purified on denaturing 8% acrylamide gels containing 8M urea. Acrylamide slices containing the RNA were eluted in water at 4°C overnight and recovered by precipitation.

### Lead Acetate Probing Assays

5’-radiolabeled *btuB* RNA (from −80 to +161 relative to the AUG) (10 nM) was incubated with increasing concentrations of Hfq (0, 1, 2.5, 5, 10 and 20 nM). Experiments were performed as previously described (69).

### Electrophoresis Mobility Shift Assays (EMSA)

For assessing the *btuB*-Hfq complex, radiolabeled *btuB* RNA (10 nM) was incubated 20 min in the absence or presence of Hfq hexamer (2.5, 5, 10, 25, 50 and 100 nM) in the protein buffer (50 mM Tris-HCl pH 7.5, 1 mM EDTA, 250 mM NH_4_Cl and 10% glycerol). For assessing the *btuB*-OmrA-Hfq complex, radiolabeled OmrA (10 nM) was incubated 10 min at 37 °C in absence of presence of Hfq (10 nM) in the protein buffer. The reaction was then incubated 20 min at 37 °C with *btuB* RNA in Afonyuskin buffer (10 mM Tris-HCl pH 7.5, 5 mM magnesium acetate, 100 mM NH_4_Cl and 0.5 mM DTT). EMSA reactions were resolved on 5% native acrylamide gels at 4 °C TBE 1X.

### Toeprinting Assays

The *btuB* RNA (0.5 pmol) was mixed with a radiolabeled DNA (3995JG; 10 nM) and incubated 5 min at 37°C. The annealing buffer was added to the reaction and incubated 5 min. When indicated, OmrA (250 nM) and/or Hfq (250 nM) were added to the mixture. Next, 30S ribosomal subunits (100 nM) and tRNA-fMet (250 nM) were added and incubated 10 min at 37 °C. The reverse transcription step was initiated by adding dNTPs (500 µM) and M-MulV-RT (10U) and the reaction was incubated 15 min at 37 °C. Reaction products were resolved on 8% denaturing acrylamide gels.

## Acknowledgements

We thank D. Schu and N. Majdalani (NIH, Bethesda) for the *hfq* variants strains, and E. Massé (Université de Sherbrooke) for strains. We are grateful to N. Fraikin and L. Van Melderen (ULB, Belgium) for the pNF02 plasmid carrying the mScarlet gene, as well as E. Hajnsdorf for the pCLHfq plasmid. We thank A. Lavigueur, M. Springer and C. Condon for critical reading of the manuscript.

This work was supported by grants from the Canadian Institutes of Health Research and the Natural Sciences and Engineering Research Council of Canada (D.A.L). This project has received funding from the European Research Council (ERC) under the European Union’s Horizon 2020 research and innovation program (Grant agreement No. 818750)(M.G.). Research in the UMR8261 is also supported by the CNRS and the “Initiative d’Excellence” program from the FrenchState (Grant “DYNAMO”, ANR-11-LABX-0011).

## References

1. Waters, L.S. and Storz, G. (2009) Regulatory RNAs in bacteria. Cell, 136, 615–628.

2. Hör, J., Matera, G., Vogel, J., Gottesman, S. and Storz, G. (2020) Trans-Acting Small RNAs and Their Effects on Gene Expression in Escherichia coli and Salmonella enterica. EcoSal Plus, 9, 10.1128/ecosalplus.ESP-0030– 2019.

3. Mccown, P.J., Corbino, K.A., Stav, S., Sherlock, M.E. and Breaker, R.R. (2017) Riboswitch diversity and distribution. RNA, 23, 995–1011.

4. Serganov, A. and Nudler, E. (2013) A decade of riboswitches. Cell, 152, 17–24.

5. Caron, M.-P., Bastet, L., Lussier, A., Simoneau-Roy, M., Massé, E. and Lafontaine, D.A. (2012) Dual-acting riboswitch control of translation initiation and mRNA decay. Proc Natl Acad Sci U S A, 109, E3444–53.

6. Bastet, L., Chauvier, A., Singh, N., Lussier, A., Lamontagne, A.M., Prévost, K., Massé, E., Wade, J.T. and Lafontaine, D.A. (2017) Translational control and Rho-dependent transcription termination are intimately linked in riboswitch regulation. Nucleic Acids Res, 45, 7474–7486.

7. Sedlyarova, N., Shamovsky, I., Bharati, B.K., Epshtein, V., Chen, J., Gottesman, S., Schroeder, R. and Nudler, E. (2016) sRNA-Mediated Control of Transcription Termination in E. coli. Cell, 167, 111–121.e13.

8. Takemoto, N., Tanaka, Y. and Inui, M. (2015) Rho and RNase play a central role in FMN riboswitch regulation in Corynebacterium glutamicum. Nucleic Acids Res, 43, 520–529.

9. Vogel, J. and Luisi, B.F. (2011) Hfq and its constellation of RNA. Nat Rev Microbiol, 9, 578–589.

10. Park, S., Prévost, K., Heideman, E.M., Carrier, M.-C., Azam, M.S., Reyer, M.A., Liu, W., Massé, E. and Fei, J. (2021) Dynamic interactions between the RNA chaperone Hfq, small regulatory RNAs, and mRNAs in live bacterial cells. Elife, 10, e64207.

11. Małecka, E.M. and Woodson, S.A. (2021) Stepwise sRNA targeting of structured bacterial mRNAs leads to abortive annealing. Mol Cell, 81, 1988–1999.

12. Holmqvist, E., Berggren, S. and Rizvanovic, A. (2020) RNA-binding activity and regulatory functions of the emerging sRNA-binding protein ProQ. Biochimica et Biophysica Acta (BBA) - Gene Regulatory Mechanisms, 1863, 194596.

13. Nou, X. and Kadner, R.J. (1998) Coupled changes in translation and transcription during cobalamin-dependent regulation of btuB expression in Escherichia coli. J Bacteriol, 180, 6719–6728.

14. Bradbeer, C., Kenley, J.S., Di Masi, D.R. and Leighton, M. (1978) Transport of vitamin B12 in Escherichia coli. Corrinoid specificities of the periplasmic B12-binding protein and of energy-dependent B12 transport. J Biol Chem, 253, 1347–1352.

15. Perdrizet 2nd, G.A., Artsimovitch, I., Furman, R., Sosnick, T.R. and Pan, T. (2012) Transcriptional pausing coordinates folding of the aptamer domain and the expression platform of a riboswitch. Proc Natl Acad Sci U S A, 109, 3323–3328.

16. Lussier, A., Bastet, L., Chauvier, A. and Lafontaine, D.A. (2015) A kissing loop is important for btuB riboswitch ligand sensing and regulatory control. J Biol Chem, 290, 26739–51.

17. Bastet, L., Chauvier, A., Singh, N., Lussier, A., Lamontagne, A.-M.M., Prévost, K., Massé, E., Wade, J.T. and Lafontaine, D.A. (2017) Translational control and Rho-dependent transcription termination are intimately linked in riboswitch regulation. Nucleic Acids Res, 45, 7474–7486.

18. Johnson Jr., J.E., Reyes, F.E., Polaski, J.T. and Batey, R.T. (2012) B12 cofactors directly stabilize an mRNA regulatory switch. Nature, 492, 133–137.

19. Guillier, M. and Gottesman, S. (2006) Remodelling of the Escherichia coli outer membrane by two small regulatory RNAs. Mol Microbiol, 59, 231–247.

20. Guillier, M. and Gottesman, S. (2008) The 5’ end of two redundant sRNAs is involved in the regulation of multiple targets, including their own regulator. Nucleic Acids Res, 36, 6781–6794.

21. De Lay, N. and Gottesman, S. (2012) A complex network of small non-coding RNAs regulate motility in Escherichia coli. Mol Microbiol, 86, 524–38.

22. Holmqvist, E., Reimegård, J., Sterk, M., Grantcharova, N., Römling, U. and Wagner, E.G.H. (2010) Two antisense RNAs target the transcriptional regulator CsgD to inhibit curli synthesis. EMBO J, 29, 1840–50.

23. Jagodnik, J., Chiaruttini, C. and Guillier, M. (2017) Stem-Loop Structures within mRNA Coding Sequences Activate Translation Initiation and Mediate Control by Small Regulatory RNAs. Mol Cell, 68, 158–170.e3.

24. Hoekzema, M., Romilly, C., Holmqvist, E. and Wagner, E.G.H. (2019) Hfq-dependent mRNA unfolding promotes sRNA-based inhibition of translation. EMBO J, 38, e101199.

25. Romilly, C., Hoekzema, M., Holmqvist, E. and Wagner, E.G.H. (2020) Small RNAs OmrA and OmrB promote class III flagellar gene expression by inhibiting the synthesis of anti-Sigma factor FlgM. RNA Biol, 17, 872– 880.

26. Brosse, A., Korobeinikova, A., Gottesman, S. and Guillier, M. (2016) Unexpected properties of sRNA promoters allow feedback control via regulation of a two-component system. Nucleic Acids Res, 44, 9650–9666.

27. Stincone, A., Daudi, N., Rahman, A.S., Antczak, P., Henderson, I., Cole, J., Johnson, M.D., Lund, P. and Falciani, F. (2011) A systems biology approach sheds new light on Escherichia coli acid resistance. Nucleic Acids Res, 39, 7512–28.

28. Kido, M., Yamanaka, K., Mitani, T., Niki, H., Ogura, T. and Hiraga, S. (1996) RNase E polypeptides lacking a carboxyl-terminal half suppress a mukB mutation in Escherichia coli. J Bacteriol, 178, 3917–3925.

29. Nahvi, A., Sudarsan, N., Ebert, M.S., Zou, X., Brown, K.L. and Breaker, R.R. (2002) Genetic control by a metabolite binding mRNA. Chem Biol, 9, 1043.

30. Franklund, C. V and Kadner, R.J. (1997) Multiple transcribed elements control expression of the Escherichia coli btuB gene. J Bacteriol, 179, 4039–4042.

31. Bandyra, K.J., Sinha, D., Syrjanen, J., Luisi, B.F. and De Lay, N.R. (2016) The ribonuclease polynucleotide phosphorylase can interact with small regulatory RNAs in both protective and degradative modes. RNA, 22, 360–372.

32. Dendooven, T., Sinha, D., Roeselová, A., Cameron, T.A., De Lay, N.R., Luisi, B.F. and Bandyra, K.J. (2021) A cooperative PNPase-Hfq-RNA carrier complex facilitates bacterial riboregulation. Mol Cell, 81, 2901–2913.e5.

33. Ravnum, S. and Andersson, D.I. (2001) An adenosyl-cobalamin (coenzyme-B12)-repressed translational enhancer in the cob mRNA of Salmonella typhimurium. Mol Microbiol, 39, 1585–1594.

34. Nou, X. and Kadner, R.J. (2000) Adenosylcobalamin inhibits ribosome binding to btuB RNA. Proc Natl Acad Sci U S A, 97, 7190–7195.

35. Kohn, H. and Widger, W. (2005) The molecular basis for the mode of action of bicyclomycin. Curr Drug Targets Infect Disord, 5, 273–95.

36. Hollands, K., Proshkin, S., Sklyarova, S., Epshtein, V., Mironov, A., Nudler, E. and Groisman, E.A. (2012) Riboswitch control of Rho-dependent transcription termination. Proc Natl Acad Sci U S A, 109, 5376–5381.

37. Figueroa-Bossi, N., Schwartz, A., Guillemardet, B., D’Heygere, F., Bossi, L. and Boudvillain, M. (2014) RNA remodeling by bacterial global regulator CsrA promotes Rho-dependent transcription termination. Genes Dev, 28, 1239–1251.

38. Mann, M., Wright, P.R. and Backofen, R. (2017) IntaRNA 2.0: enhanced and customizable prediction of RNA-RNA interactions. Nucleic Acids Res, 45, W435–W439.

39. Jagodnik, J., Chiaruttini, C. and Guillier, M. (2017) Stem-Loop Structures within mRNA Coding Sequences Activate Translation Initiation and Mediate Control by Small Regulatory RNAs. Mol Cell, 68, 158–170.e3.

40. Quinn, H.J., Cameron, A.D.S. and Dorman, C.J. (2014) Bacterial regulon evolution: distinct responses and roles for the identical OmpR proteins of Salmonella Typhimurium and Escherichia coli in the acid stress response. PLoS Genet, 10, e1004215.

41. Schu, D.J., Zhang, A., Gottesman, S. and Storz, G. (2015) Alternative Hfq-sRNA interaction modes dictate alternative mRNA recognition. EMBO J, 34, 2557–73.

42. Zhang, A., Schu, D.J., Tjaden, B.C., Storz, G. and Gottesman, S. (2013) Mutations in interaction surfaces differentially impact E. coli Hfq association with small RNAs and their mRNA targets. J Mol Biol, 425, 3678– 97.

43. Desnoyers, G. and Masse, E. (2012) Noncanonical repression of translation initiation through small RNA recruitment of the RNA chaperone Hfq. Genes Dev, 26, 726–739.

44. Azam, M.S. and Vanderpool, C.K. (2018) Translational regulation by bacterial small RNAs via an unusual Hfq-dependent mechanism. Nucleic Acids Res, 46, 2585–2599.

45. Ziolkowska, K., Derreumaux, P., Folichon, M., Pellegrini, O., Régnier, P., Boni, I. V and Hajnsdorf, E. (2006) Hfq variant with altered RNA binding functions. Nucleic Acids Res, 34, 709–20.

46. Hartz, D., McPheeters, D.S., Traut, R. and Gold, L. (1988) Extension inhibition analysis of translation initiation complexes. Methods Enzymol, 164, 419–25.

47. Kashket, E.R. (1981) Proton motive force in growing Streptococcus lactis and Staphylococcus aureus cells under aerobic and anaerobic conditions. J Bacteriol, 146, 369–76.

48. Minamino, T., Imae, Y., Oosawa, F., Kobayashi, Y. and Oosawa, K. (2003) Effect of intracellular pH on rotational speed of bacterial flagellar motors. J Bacteriol, 185, 1190–4.

49. Bradbeer, C. (1993) The proton motive force drives the outer membrane transport of cobalamin in Escherichia coli. J Bacteriol, 175, 3146–50.

50. Lei, G.-S., Syu, W.-J., Liang, P.-H., Chak, K.-F., Hu, W.S. and Hu, S.-T. (2011) Repression of btuB gene transcription in Escherichia coli by the GadX protein. BMC Microbiol, 11, 33.

51. Storz, G., Vogel, J. and Wassarman, K.M. (2011) Regulation by small RNAs in bacteria: expanding frontiers. Mol Cell, 43, 880–91.

52. Wagner, E.G.H. and Romby, P. (2015) Small RNAs in bacteria and archaea: who they are, what they do, and how they do it. Adv Genet, 90, 133–208.

53. Nitzan, M., Rehani, R. and Margalit, H. (2017) Integration of Bacterial Small RNAs in Regulatory Networks. Annu Rev Biophys, 46, 131–148.

54. Reyer, M.A., Chennakesavalu, S., Heideman, E.M., Ma, X., Bujnowska, M., Hong, L., Dinner, A.R., Vanderpool, C.K. and Fei, J. (2021) Kinetic modeling reveals additional regulation at co-transcriptional level by post-transcriptional sRNA regulators. Cell Rep, 36, 109764.

55. Chen, J., Morita, T. and Gottesman, S. (2019) Regulation of Transcription Termination of Small RNAs and by Small RNAs: Molecular Mechanisms and Biological Functions. Front Cell Infect Microbiol, 9, 201.

56. Bossi, L., Figueroa-Bossi, N., Bouloc, P. and Boudvillain, M. (2020) Regulatory interplay between small RNAs and transcription termination factor Rho. Biochim Biophys Acta Gene Regul Mech, 1863, 194546.

57. Bossi, L., Schwartz, A., Guillemardet, B., Boudvillain, M. and Figueroa-Bossi, N. (2012) A role for Rho-dependent polarity in gene regulation by a noncoding small RNA. Genes Dev, 26, 1864–1873.

58. Wickiser, J.K., Winkler, W.C., Breaker, R.R. and Crothers, D.M. (2005) The speed of RNA transcription and metabolite binding kinetics operate an FMN riboswitch. Mol Cell, 18, 49–60.

59. Lemay, J.-F., Desnoyers, G., Blouin, S., Heppell, B., Bastet, L., St-Pierre, P., Massé, E. and Lafontaine, D.A. (2011) Comparative study between transcriptionally- and translationally-acting adenine riboswitches reveals key differences in riboswitch regulatory mechanisms. PLoS Genet, 7, e1001278.

60. Morita, T., Maki, K. and Aiba, H. (2005) RNase E-based ribonucleoprotein complexes: mechanical basis of mRNA destabilization mediated by bacterial noncoding RNAs. Genes Dev, 19, 2176–2186.

61. Caillet, J., Baron, B., Boni, I. V, Caillet-Saguy, C. and Hajnsdorf, E. (2019) Identification of protein-protein and ribonucleoprotein complexes containing Hfq. Sci Rep, 9, 14054.

62. Loh, E., Dussurget, O., Gripenland, J., Vaitkevicius, K., Tiensuu, T., Mandin, P., Repoila, F., Buchrieser, C., Cossart, P. and Johansson, J. (2009) A trans-acting riboswitch controls expression of the virulence regulator PrfA in Listeria monocytogenes. Cell, 139, 770–779.

63. DebRoy, S., Gebbie, M., Ramesh, A., Goodson, J.R., Cruz, M.R., van Hoof, A., Winkler, W.C. and Garsin, D. a. (2014) Riboswitches. A riboswitch-containing sRNA controls gene expression by sequestration of a response regulator. Science, 345, 937–940.

64. Mellin, J.R., Koutero, M., Dar, D., Nahori, M.-a. A., Sorek, R. and Cossart, P. (2014) Riboswitches. Sequestration of a two-component response regulator by a riboswitch-regulated noncoding RNA. Science, 345, 940–943.

65. Melamed, S., Adams, P.P., Zhang, A., Zhang, H. and Storz, G. (2020) RNA-RNA Interactomes of ProQ and Hfq Reveal Overlapping and Competing Roles. Mol Cell, 77.

66. Miller, J.H. (1972) Experiments in Molecular Genetics Laboratory, C.S.H. (ed) Cold Spring Harbor Laboratory Press, NY, USA.

67. Lemay, J.F., Penedo, J.C., Tremblay, R., Lilley, D.M.J. and Lafontaine, D.A.A. (2006) Folding of the Adenine Riboswitch. Chem Biol, 13, 857–868.

68. Aiba, H., Adhya, S. and de Crombrugghe, B. (1981) Evidence for two functional gal promoters in intact Escherichia coli cells. J Biol Chem, 256, 11905–11910.

69. Desnoyers, G., Morissette, A., Prevost, K. and Masse, E. (2009) Small RNA-induced differential degradation of the polycistronic mRNA iscRSUA. EMBO J, 28, 1551–1561.

