## Supplementary Materials for "Riboswitch and small RNA modulate *btuB* translation initiation in *Escherichia coli* and trigger distinct mRNA regulatory mechanisms"

**Supplementary Information**

**Strains and plasmids constructions**

Strains and plasmids constructed and used in this study are listed in the Supplementary Table S1. Corresponding DNA oligonucleotides (oligos) are summarized in the Supplementary Table S2.

Genetic manipulations for the construction of new gene knockouts and reporter fusions were carried out using a recombineering approach that employs phage λ Red recombination functions as described in detail in (1). Simple allelic exchange was performed using the generalized P1 transduction as described in (2). Strains were cultivated in liquid LB, LB agar plates, BYE (LB without salt) agar plates supplemented with 6% (w/v) sucrose (BS plates) or liquid and solid minimal medium A. Acid LB (pH 4.7) was made by addition of HCl. When needed, media were supplemented with an appropriate antibiotic. Unless otherwise stated, kanamycin (Kan) at 50 µg/ml, chloramphenicol (Cam) at 20 µg/ml, ampicillin (Amp) at 150 µg/ml or tetracycline (Tet) at 10 µg/ml were utilized for plasmid maintenance. For selection of strains carrying a single copy of a drug resistance marker the concentrations were twice as low. During the P1 transduction, selectable plates also contained 5 mM sodium citrate.

*Designing an E. coli strain for construction of chromosomal mScarlet fluorescent reporter fusions*

The strain OK510 was engineered for constructing mScarlet fusions. It is a DJ624 (MG1655 *ΔlacX74 mal::lacIq*) derivative that carries a *mini-λ-Tet* prophage (3), which provides λ Red recombination functions that are induced at 42°C, and an *mScarlet* locus placed into the 6 bp intergenic region *argG-yhbX*. This locus (see Supplementary Figure S11) comprises a synthetic bidirectional transcriptional terminator L3S2P21 (TT1) (4), an inactive $P_{LtetO-1}$ promoter (5) that is deprived of the -10 region, the Pcat-*cat-sacB* cassette expressed in the opposite direction to that of following 'mScarlet region (mScarlet-I ORF missing the translation initiation codon), and the *nptII* ORF, which is transcriptionally coupled to *'mScarlet* and flanked by two FRT sites. The FRT sites allow for optional elimination of the *nptII* ORF *via* Flp-FRT recombination. The last element of the locus is a natural bidirectional transcriptional terminator, ECK120026481 (TT2) (4). Hence, the strain features $Tet^R$ (due to the presence of *mini-λ-Tet*), $Sucr^S$, $Cam^R$ and $Kan^S$ as both '*mScarlet* and *nptII* genes are not expressed.

The construction of mScarlet fusions was made with PCR cassette containing ~40 bp homologies on both ends that allow for recombining with its 5'-end within the inactive $P_{LtetO-1}$ restoring the -10 region, and with its 3'-end within the '*mScarlet* (restoring the translation initiation codon), so that it replaces the Pcat-*cat-sacB* sequences. This generates a translational fusion to *mScarlet*, which is expressed from a strong constitutive (in the absence of the Tet repressor) promoter $P_{LtetO-1}$ and provides expression of the downstream *nptII* ORF that renders the recombinants resistance to kanamycin. Thus, the recombinants can be selected for both resistance to sucrose and kanamycin, which greatly reduces occurrence of false-positive clones as compared to selection on sucrose alone. Selection at 37°C also facilitates elimination the *mini-λ-Tet* prophage in the majority of recombinants.

This OK510 strain was made through multiple steps of recombineering, summarized in Supplementary Figure S11. First, a *cat-sacB* cassette was amplified in 3 steps in order to add upstream the inactive $P_{LtetO-1}$ promoter lacking the -10 region, as well as upstream and downstream transcription terminators (TT1 and TT2, respectively) and homology regions to the argG/yhbX locus. The first step PCR template was the genomic DNA of a strain carrying a *cat-sacB* cassette and the primers used are (i) Ptetno-10-cat-for and Ter-catsacRev (1st step), (ii) Ter-Ptet-for and yhbX-Ter-rev (2nd step = reamplification of PCR of 1st step) and (iii) argG-Ter-for and yhbX-Ter-rev (3rd step = reamplification of PCR of 2nd step). This last PCR product was recombined in strain MG1432 and recombinants were selected on LB-Cam at 30°C, and then checked for sucrose sensitivity and Tet resistance (to ensure that they kept the mini-λ Tet for further recombineering). The structure of the locus was checked by sequencing and the resulting strain was called MG2346. In a second recombineering step, a *nptII* gene followed by an FRT site (PCR product amplified from genomic DNA with primers sacB-KanR-For and FRT-sacB-Rev) was introduced downstream of the *sacB* gene, giving rise to the strain MG2348 after selection on Kan and sequencing of the locus. The third step consisted in replacing the *cat-sacB* cassette by the mScarlet gene devoid of the start codon, and followed by an FRT site. This was again done by recombineering, into strain MG2348 this time, using a PCR product amplified from the pNF02-mScI plasmid (a gift from N. Fraikin and L. Van Melderen, (6)) with primers Ptetno-10-mSC-For and mSc-FRT-Rev, and then reamplified with primers Ptet-55-12For and postFRT-KanR-Rev to increase the length of the homology regions. Recombinants were selected at 30°C on BS plates, and checked for CamS and TetR. Clones whose structure of this TT1-Ptet(no-10)-‘mScarlet(no AUG)-FRT-nptII-FRT-TT2 region was confirmed by sequencing were found to be KanS; out of those, strain MG2352 was used for the subsequent step.

At the last step, the Pcat-*cat-sacB* cassette, made with oligos AK411 and AK412 using chromosomal DNA of NC397 strain (7) as a template, was recombined into MG2352 with subsequent selection of recombinants on LB-Cam plates at 30°C giving rise to the strain OK509. To avoid mutations that could have accumulated upon multiple recombineering passages during construction of the OK509 strain, the ‘*mScarlet* locus of OK509 was transferred into the initial strain, MG1432, selecting transductants on LB-Cam plates at 30°C. The resulting strain, OK510, was used for construction of the mScarlet fusions used in this study.

*mScarlet fusions construction*

To obtain $P_{LtetO-1}$-*sdhC*$_{-222+39}$-*mScarlet*$_{+4}$, $P_{LtetO-1}$-*ompR*$_{-35+717}$-*mScarlet*$_{+4}$, and $P_{LtetO-1}$-*btuB*$_{-240+210}$-*mScarlet*$_{+4}$, corresponding *loci* of MG1655 chromosome were PCR amplified with following pairs of oligos: Ptet-sdhC-222for and sdhC+39-mScrev (then, re-amplified with Ptet-55-12For and sdhC+39-mScrev); 5'PtetompR-35+30-lacZ and ompR+717-mScrev (then, re-amplified with Ptet-55-12For and ompR+717-mScrev); 5’PtetBtuB-240 and AK451 (then, re-amplified with Ptet-55-12For and AK420). Resulting PCR products were recombined into the OK510 strain. Recombinants were selected at 37°C on BS plates supplemented with 6 µg/ml kanamycin. The recombinant colonies were checked for fluorescence and sensitivity to tetracycline (elimination of the mini-λ-Tet). The upstream *mScarlet* regions of the obtained OK528, OK529 and OK572 strains were sequenced with oligos AK387 and AK418.

*lacZ fusions construction*

Construction of *lacZ* fusions was done by replacing a *cat-sacB* cassette upstream of *lacZ* by recombineering into strain PM1205 ((8), for construction of $P_{BAD}$ driven fusions) or MG1508 ((9), for construction of $P_{LtetO-1}$ driven fusions). In general, the PCR products were obtained using MG1655 genomic DNA as a template, and, when needed, the length of the homology regions was extended by re-amplification with primers Ptet-55-12For or lacZ28-66rev as necessary. The mutations EP1, EP2, EP3, M9, M11 and mutH1 were present on the primers, and the same was true for the low$P_{LtetO-1}$ mutant that changes the -35 region consensus of $P_{LtetO-1}$ from TTGACA into TGGACA. For the construction of the $P_{BAD}$-*btuB-lacZ* transcriptional fusions, successive PCR reactions introduced, downstream of the selected *btuB* region, an in-frame sequence encoding a DPAF peptide terminated by a stop codon, followed by *lacZ* sequence starting 17nts before *lacZ* translation initiation codon.

Recombinants were selected on BS plates supplemented with 0,002% X-gal at 37°C, picking blue colonies and checking them for sensitivity to chloramphenicol (loss of the *cat-sac* cassette) and tetracycline (loss of the *mini-λ-Tet*). The *lac* loci of resulting strains were verified by sequencing.

*Constructing omrA and omrB and omrAB deletion strains*

PCR cassettes for generating *ΔomrA::nptI* and *ΔomrB::nptI* were obtained using pairs of primers, OmrA-kan5 + OmrA-Kan3 and OmrB-kan5 + OmrB-kan3, respectively. As a template, chromosomal DNA of a strain carrying an *nptI* gene was used. The cassettes were recombined into NM300 strain selecting recombinants on LB-Kan at 37°C. Loss of mini-λ-Tet was verified by sensitivity to Tet. The deletions in resulting strains MG1001 (*ΔomrA::nptI*) and MG1002 (*ΔomrB::nptI*) were verified by PCR and sequenced with oligos seqOmrBfor and seqOmrArev. The obtained *ΔomrA::nptI* and *ΔomrB::nptI* and the previously published double deletion *ΔomrAB::nptI* of MG1003 (10) were transferred to

JJ416, generating OK615, OK616 and JJ426, respectively. Same deletions were transferred to JJ425 giving rise to OK617, OK618, and JJ427, respectively.

*Elimination of the nptII ORF from FRT-nptII-FRT cassettes*

Unnecessary *nptII* ORFs originating from the OK510 derivatives and from Keio collection knockouts (11) used in this study were eliminated *via* Flp-FRT recombination. To this end, cells were transformed with the pCP20 plasmid (12). The plasmid provides *in trans* synthesis of flippase (Flp) and possesses a thermo-sensitive replication origin. Transformants were selected at 30°C on LB-Cam (10 μg/ml) plates supplemented with chloramphenicol and purified once on the same medium. Then, an individual transformant colony was grown overnight in 10 ml of LB medium at 42°C and plated on LB at ~$10^2$ cfu per plate 37°C. After such passage vast majority of clones were both KanS and CamS.

*Constructing Hfq variant strains carrying the mScarlet fusions under study*

The construction of strains carrying different *hfq* mutations is based on (13), with minor modifications to combine these different alleles with mScarlet fusions. First, the *ΔargG*::FRT-*nptII*-FRT allele was transferred from JW3140 (from the Keio collection (11)) to DJ624 by P1 transduction to generate strain OK523. This OK523 strain was then cured of *nptII* as described above generating OK530. The *Δhfq::cat-sacB* allele was co-transduced with nearby *purA::*FRT-*nptII*-FRT from DJS2604 (from D. Schu, NIH) to OK530 selecting on LB-Cam, thus giving rise to OK564. The *hfq* alleles were transferred from DJS2609 (*hfq* WT), DJS2927 (*Δhfq*), KK2560 (*hfqQ8A*), KK2561 (*hfqR16A*), and KK2562 (*hfqY25D*) to OK564 selecting transductants on minimal medium A-agar plates (14) supplemented with 0.2% (w/v) of glucose and 20 μg/ml of L-arginine (the acceptor strain, OK564, is a purine and arginine auxotroph). Transfer of the *hfq* alleles was verified by PCR with *hfq* check oligos mHfqout and antiHfqout, or with the mHfqout and one of the *hfq* point mutant-specific oligos, AK430 (Q8A-specific), AK431 (R16A-specific), and AK432 (Y25D-specific). The resulting strains, OK581 (*hfq* WT), OK582 (*Δhfq*), OK583 (*hfqQ8A*), OK584 (*hfqR16A*), and OK585 (*hfqY25D*), served as acceptors for transferring the mScarlet fusion variants. To do so, the fusion strains OK528, OK529 and OK572 were cured of the *nptII* cassette, thus generating strains OK560, OK561 and OK577, respectively. Then, each fusion was transduced into the five *hfq* variant strains. Arginine prototroph transductants were selected at 37°C on minimal A-agar plates supplemented with 0.2% glucose (mScarlet locus is 100% co-transduced with the wild type *argG* allele). The resulting strains, OK586 to OK605, are listed in the Supplementary Table S1.

*Fluorescence measurements to assess the expression of mScarlet fusions*

The *hfq* variant strains OK586-OK590 ($P_{LtetO\text{-}1}$-*btuB*$_{-240+210}$-*mScarlet*$_{+4}$), and OK601-OK605 ($P_{LtetO\text{-}1}$-*ompR*$_{-35+717}$-*mScarlet*$_{+4}$) were transformed with the pBRplac vector control, pOmrA and pOmrB. The $P_{LtetO\text{-}1}$-*sdhC*$_{-222+39}$-*mScarlet*$_{+4}$ *hfq* variant strains (OK596-OK600) were transformed with pBRplac and pSpot42. In each case transformants were selected on LB-Tet plates at 37°C. DJ624 transformed with pBRplac was utilized as a no-fusion background. After a single purification on the LB-Tet plate, individual transformant colony was inoculated with 400 µl of CAG medium (minimal A salts (14), 0.5% (w/v) glycerol, 0.25% (w/v) casamino acids, 1 mM $MgSO_4$) supplemented with Tet and incubated overnight with shaking at 37°C. Next day, 0.4 µl of saturated culture used to inoculate 200 µl of CAG medium supplemented with Tet and 0.25 mM IPTG in the black 96 well µCLEAR F-bottom plate (Greiner, 655090) covered with 50 µl of mineral oil (Sigma, M8410).

Test was run in the CLARIOStar$^{PLUS}$ plate reader (BMG Labtech) at 37°C and 500 RPM. Absorbance at 600 nm and fluorescence (excitation at 560±15 nm, emission at 600±15 nm with 580 nm dichroic filter) was measured every 12 minutes for 16 hours. Each experiment was made in triplicate (starting from three independent transformant colonies). Experimental curves expressed as fluorescence normalized to absorbance at 600 nm (normalized fluorescence) versus absorbance at 600 nm are shown in the Supplementary Figures S2 and S4-S9. To generate the histograms, the mean normalized fluorescence and standard deviations were calculated for the three (one per triplicate) time points closest to apparent $A_{600}$=0.3. Mean normalized fluorescence of the non-fusion control was calculated accordingly and subtracted from the experimental data.

#### References

1. Sawitzke,J.A., Thomason,L.C., Costantino,N., Bubunenko,M., Datta,S. and Court,D.L. (2007) Recombineering: in vivo genetic engineering in E. coli, S. enterica, and beyond. *Methods Enzymol*, **421**, 171–99.
2. Thomason,L.C., Costantino,N. and Court,D.L. (2007) E. coli genome manipulation by P1 transduction. *Curr Protoc Mol Biol*, **1**, 1.17.1-1.17.8.
3. Court,D.L., Swaminathan,S., Yu,D., Wilson,H., Baker,T., Bubunenko,M., Sawitzke,J. and Sharan,S.K. (2003) Mini-lambda: a tractable system for chromosome and BAC engineering. *Gene*, **315**, 63–9.
4. Chen,Y.-J., Liu,P., Nielsen,A.A.K., Brophy,J.A.N., Clancy,K., Peterson,T. and Voigt,C.A. (2013) Characterization of 582 natural and synthetic terminators and quantification of their design constraints. *Nat Methods*, **10**, 659–64.
5. Lutz,R. and Bujard,H. (1997) Independent and tight regulation of transcriptional units in Escherichia coli via the LacR/O, the TetR/O and AraC/I1-I2 regulatory elements. *Nucleic Acids Res*, **25**, 1203–10.
6. Fraikin,N., Rousseau,C.J., Goeders,N. and Van Melderen,L. (2019) Reassessing the Role of the Type II MqsRA Toxin-Antitoxin System in Stress Response and Biofilm Formation: mqsA Is Transcriptionally Uncoupled from mqsR. *mBio*, **10**, e0267819.
7. Svenningsen,S.L., Costantino,N., Court,D.L. and Adhya,S. (2005) On the role of Cro in lambda prophage induction. *Proc Natl Acad Sci U S A*, **102**, 4465–9.
8. Mandin,P. and Gottesman,S. (2009) A genetic approach for finding small RNAs regulators of genes of interest identifies RybC as regulating the DpiA/DpiB two-component system. *Mol Microbiol*, **72**, 551–565.
9. Coornaert,A., Chiaruttini,C., Springer,M. and Guillier,M. (2013) Post-transcriptional control of the Escherichia coli PhoQ-PhoP two-component system by multiple sRNAs involves a novel pairing region of GcvB. *PLoS Genet*, **9**, e1003156.
10. Guillier,M. and Gottesman,S. (2006) Remodelling of the Escherichia coli outer membrane by two small regulatory RNAs. *Mol Microbiol*, **59**, 231–247.
11. Baba,T., Ara,T., Hasegawa,M., Takai,Y., Okumura,Y., Baba,M., Datsenko,K.A., Tomita,M., Wanner,B.L. and Mori,H. (2006) Construction of Escherichia coli K-12 in-frame, single-gene knockout mutants: the Keio collection. *Mol Syst Biol*, **2**, 2006.0008.
12. Cherepanov,P.P. and Wackernagel,W. (1995) Gene disruption in Escherichia coli: TcR and KmR cassettes with the option of Flp-catalyzed excision of the antibiotic-resistance determinant. *Gene*, **158**, 9–14.
13. Zhang,A., Schu,D.J., Tjaden,B.C., Storz,G. and Gottesman,S. (2013) Mutations in interaction surfaces differentially impact E. coli Hfq association with small RNAs and their mRNA targets. *J Mol Biol*, **425**, 3678–97.
14. Miller,J.H. (1972) Experiments in Molecular Genetics Laboratory,C.S.H. (ed) Cold Spring Harbor Laboratory Press, NY, USA.
15. Madeira,F., Park,Y.M., Lee,J., Buso,N., Gur,T., Madhusoodanan,N., Basutkar,P., Tivey,A.R.N., Potter,S.C., Finn,R.D., *et al.* (2019) The EMBL-EBI search and sequence analysis tools APIs in 2019. *Nucleic Acids Res*, **47**, W636–W641.

16. Zuker,M. (2003) Mfold web server for nucleic acid folding and hybridization prediction. *Nucleic Acids Res*, **31**, 3406–3415.
17. Masse,E. and Gottesman,S. (2002) A small RNA regulates the expression of genes involved in iron metabolism in Escherichia coli. *Proc Natl Acad Sci U S A*, **99**, 4620–4625.
18. Masse,E., Escorcia,F.E. and Gottesman,S. (2003) Coupled degradation of a small regulatory RNA and its mRNA targets in Escherichia coli. *Genes Dev*, **17**, 2374–2383.
19. Bastet,L., Chauvier,A., Singh,N., Lussier,A., Lamontagne,A.M., Prévost,K., Massé,E., Wade,J.T. and Lafontaine,D.A. (2017) Translational control and Rho-dependent transcription termination are intimately linked in riboswitch regulation. *Nucleic Acids Res*, **45**, 7474–7486.
20. Jagodnik,J., Chiaruttini,C. and Guillier,M. (2017) Stem-Loop Structures within mRNA Coding Sequences Activate Translation Initiation and Mediate Control by Small Regulatory RNAs. *Mol Cell*, **68**, 158-170.e3.
21. Majdalani,N., Cunning,C., Sledjeski,D., Elliott,T. and Gottesman,S. (1998) DsrA RNA regulates translation of RpoS message by an anti-antisense mechanism, independent of its action as an antisilencer of transcription. *Proc Natl Acad Sci U S A*, **95**, 12462–12467.
22. Mandin,P. and Gottesman,S. (2010) Integrating anaerobic/aerobic sensing and the general stress response through the ArcZ small RNA. *EMBO J*, **29**, 3094–107.
23. Lerner,C.G. and Inouye,M. (1990) Low copy number plasmids for regulated low-level expression of cloned genes in Escherichia coli with blue/white insert screening capability. *Nucleic Acids Res*, **18**, 4631.
24. Ziolkowska,K., Derreumaux,P., Folichon,M., Pellegrini,O., Régnier,P., Boni,I. V and Hajnsdorf,E. (2006) Hfq variant with altered RNA binding functions. *Nucleic Acids Res*, **34**, 709–20.
25. Bindels,D.S., Haarbosch,L., van Weeren,L., Postma,M., Wiese,K.E., Mastop,M., Aumonier,S., Gotthard,G., Royant,A., Hink,M.A., *et al.* (2017) mScarlet: a bright monomeric red fluorescent protein for cellular imaging. *Nat Methods*, **14**, 53–56.

**A**

```
OmrA CCCAGAGGUAUUGAUUGGUGAGAUUAU--UCGGUACGCUCUUCGUACCCUGUCUCUUGCA 58
OmrB CCCAGAGGUAUUGAUAGGUGAAGUCAACUUCGGGU-----U---GAGCACAUGAAUUACA 52
     *************** *****  * *   ****       *    * *   *   ** **

OmrA CCAACCUGCGCGGAUGCGCAGGUUUUUUUU 88
OmrB CCAGCCUGCGCAGAUGCGCAGGUUUUUUUU 82
     *** ******* ******************
```

**B**

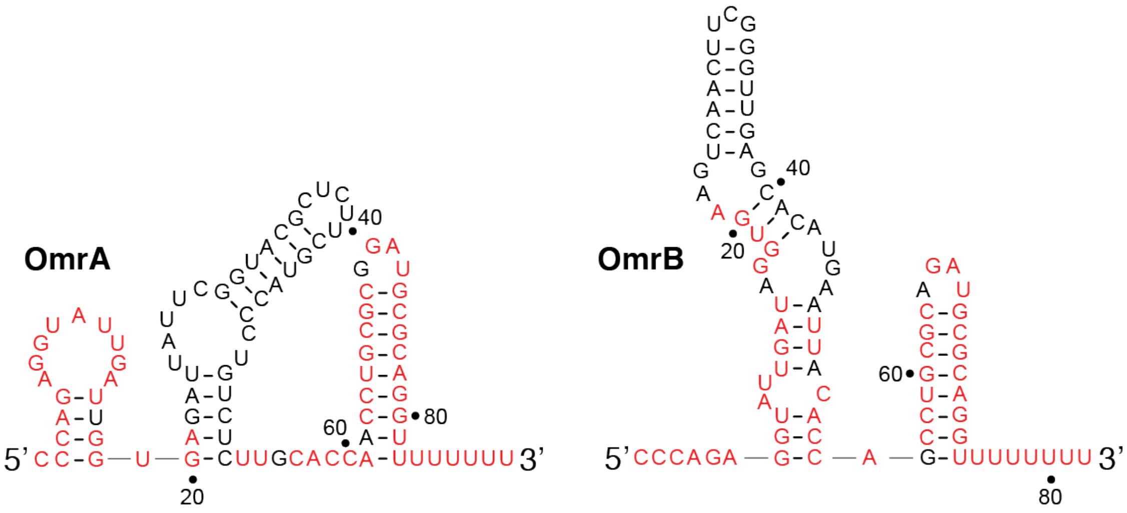

**Supplementary Figure S1. Sequences and secondary structures of OmrA and OmrB.**

**(A)** Sequence alignment of *E. coli* OmrA and OmrB sRNAs performed using Clustal (15). Nucleotides in red represent the 5' and 3' conserved regions of sRNAs. **(B)** Secondary structures of OmrA and OmrB. The structures have been predicted using Mfold (16). Nucleotides in red indicate the 5' and 3' conserved regions of both sRNAs.

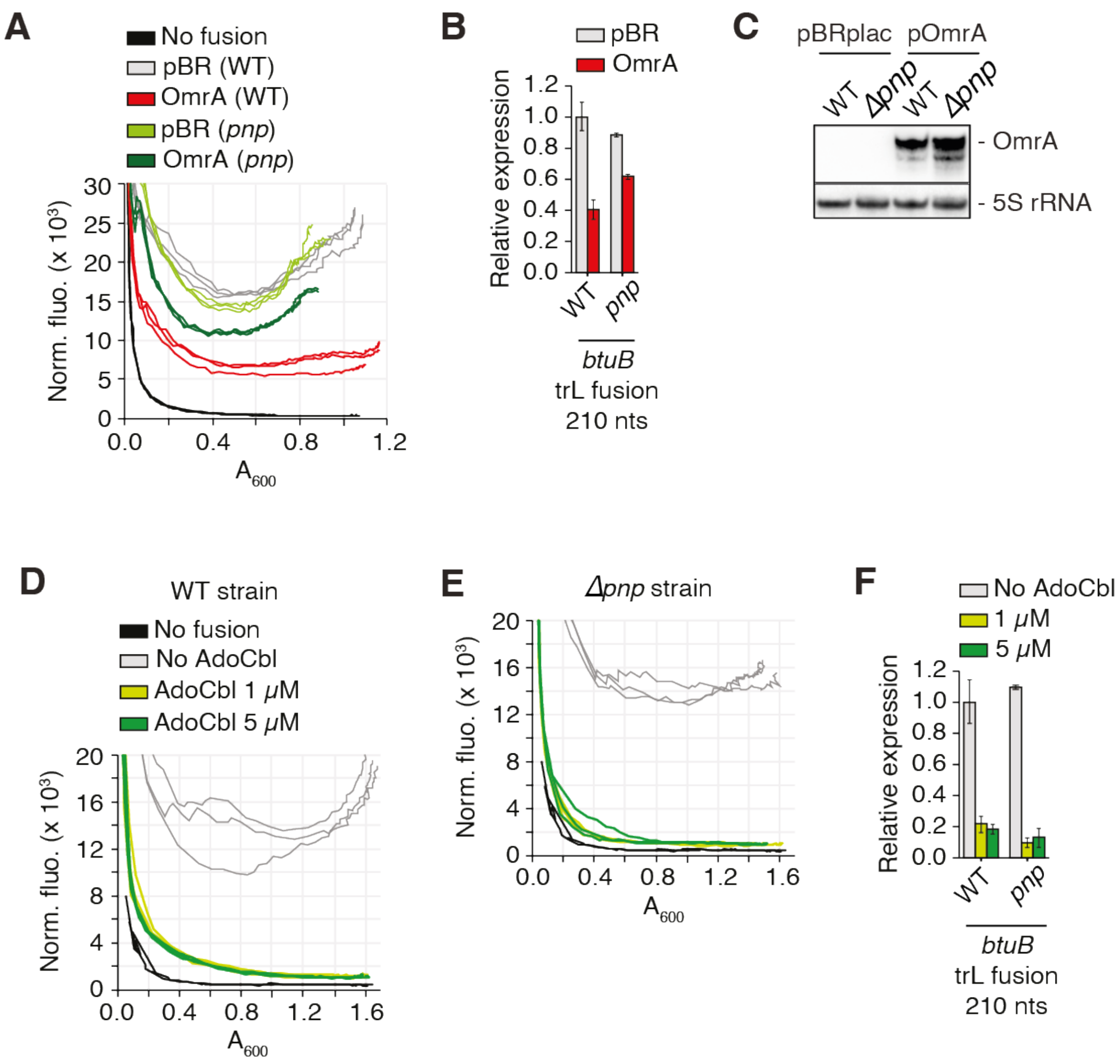

**Supplementary Figure S2. The control of *btuB* by OmrA is impaired in the absence of PNPase. (A)** The fluorescence of the $BtuB_{210}$-mScarlet translational fusion was followed in WT and *pnp* deleted cells, transformed with a vector control or an OmrA overproducing plasmid. The fluorescence of a strain that does not express the mScarlet gene was followed as a control for the fluorescence background. The curves show normalized fluorescence plotted against the absorbance at 600nm for all samples (in triplicate). **(B)** The average value and standard deviation of normalized fluorescence at an absorbance of 600nm closest to 0.3 is shown on the histograms. **(C)** The levels of OmrA in WT or *pnp* deleted cells were analyzed by Northern-blot in an independent experiment. **(D, E, F)** The fluorescence of the $BtuB_{210}$-mScarlet fusion was measured in WT (D) or *pnp* deleted cells (E) grown in the absence or in the presence of AdoCbl at a final concentration of 1 or 5 µM. Graphic representation is as in panel A, and data from panels D and E are processed and summarized on panel F the same way as for panel B.

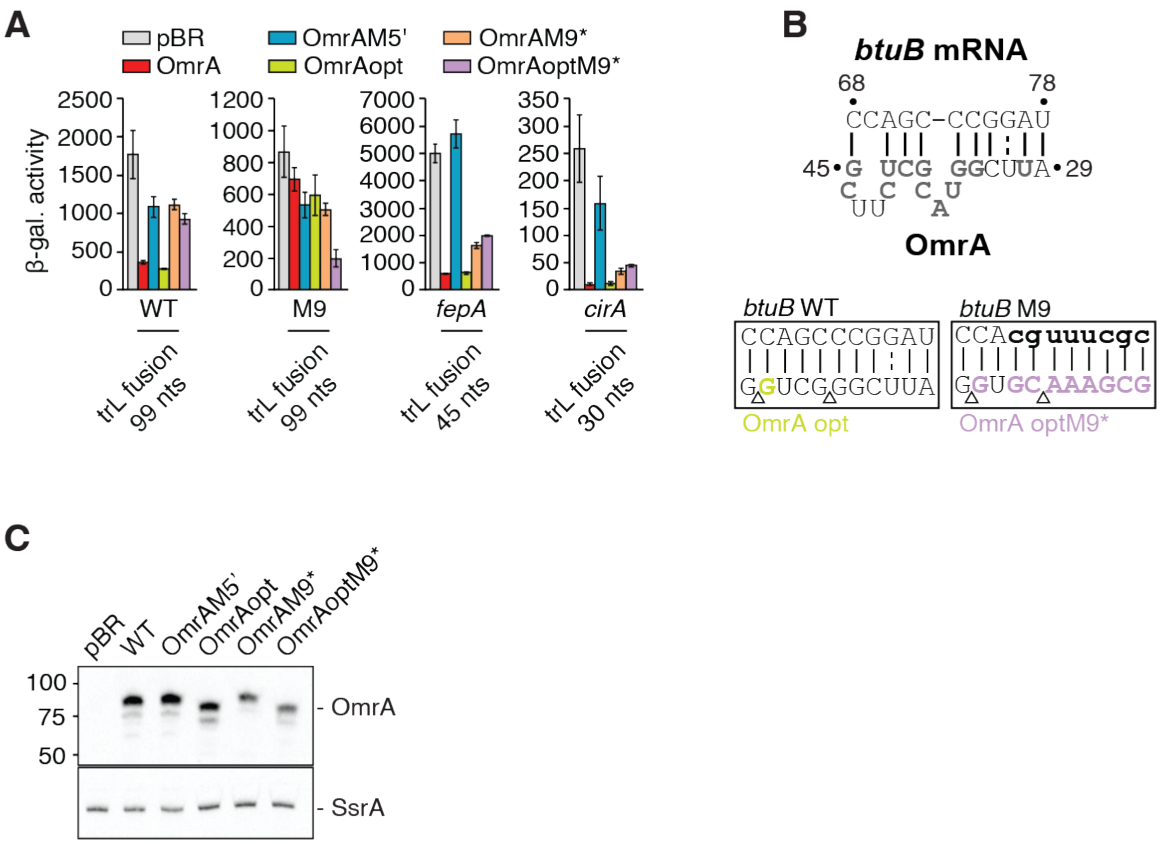

**Supplementary Figure S3. Sequence changes at the 5' end and central region of OmrA modulates *btuB* regulation. (A)** Beta-galactosidase assays of $BtuB_{99}$-LacZ (WT and M9 mutant, $FepA_{45}$-LacZ and $CirA_{30}$-LacZ translational fusions upon overproduction of different OmrA variants. The b-galactosidase average values (in Miller units) and the standard deviations were obtained from three independent experiments. All fusions are expressed from a $P_{LtetO-1}$ constitutively expressed promoter. **(B)** Predicted OmrA-*btuB* interactions are shown for key sRNA-mRNA pairs for the WT and OmrAopt, as well as for *btuB* M9 mutant and OmrAoptM9*. Lowercase and colored nucleotides represent mutations introduced in *btuB* M9 and OmrA, respectively; deletion of nts in OmrAopt is indicated by the triangles. **(C)** Northern blot analysis of levels of OmrA variants using

RNA extracted from cell cultures used for corresponding ß-galactosidase assays. Detection of the SsrA RNA was used as a loading control.

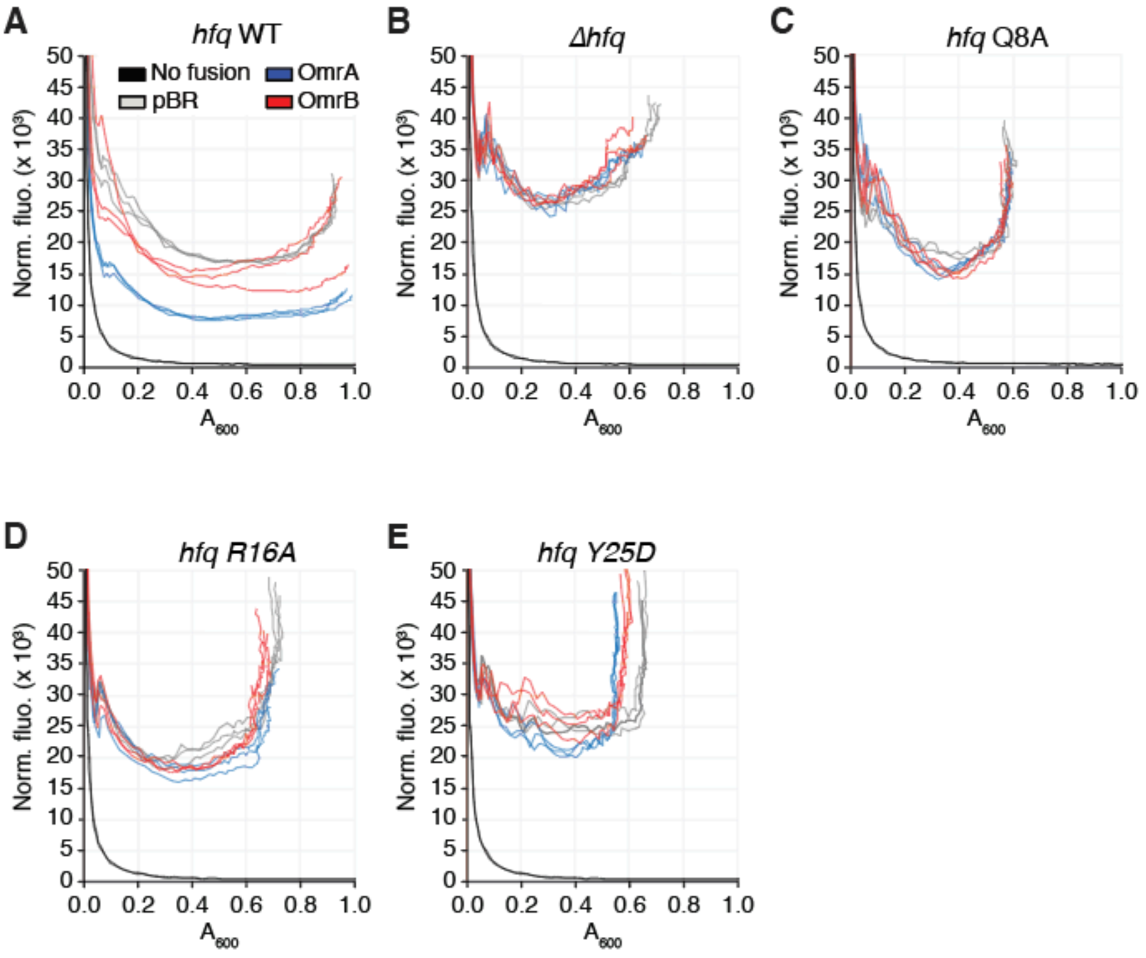

**Supplementary Figure S4. Fluorescence assays using the $BtuB_{210}$-mScarlet fusion. (A-E)** Experiments were performed when overexpressing OmrA, OmrB or when using the empty vector (pBR). A control was performed without the fluorescent construct (no fusion). Experiments were performed in the WT strain (*hfq* WT) **(A)**, Δ*hfq* **(B)**, *hfq* Q8A **(C)**, *hfq* R16A **(D)** and *hfq* Y25D **(E)**. These raw data were used to calculate the relative FPA (fluorescence per absorbance) shown in Figure 4B.

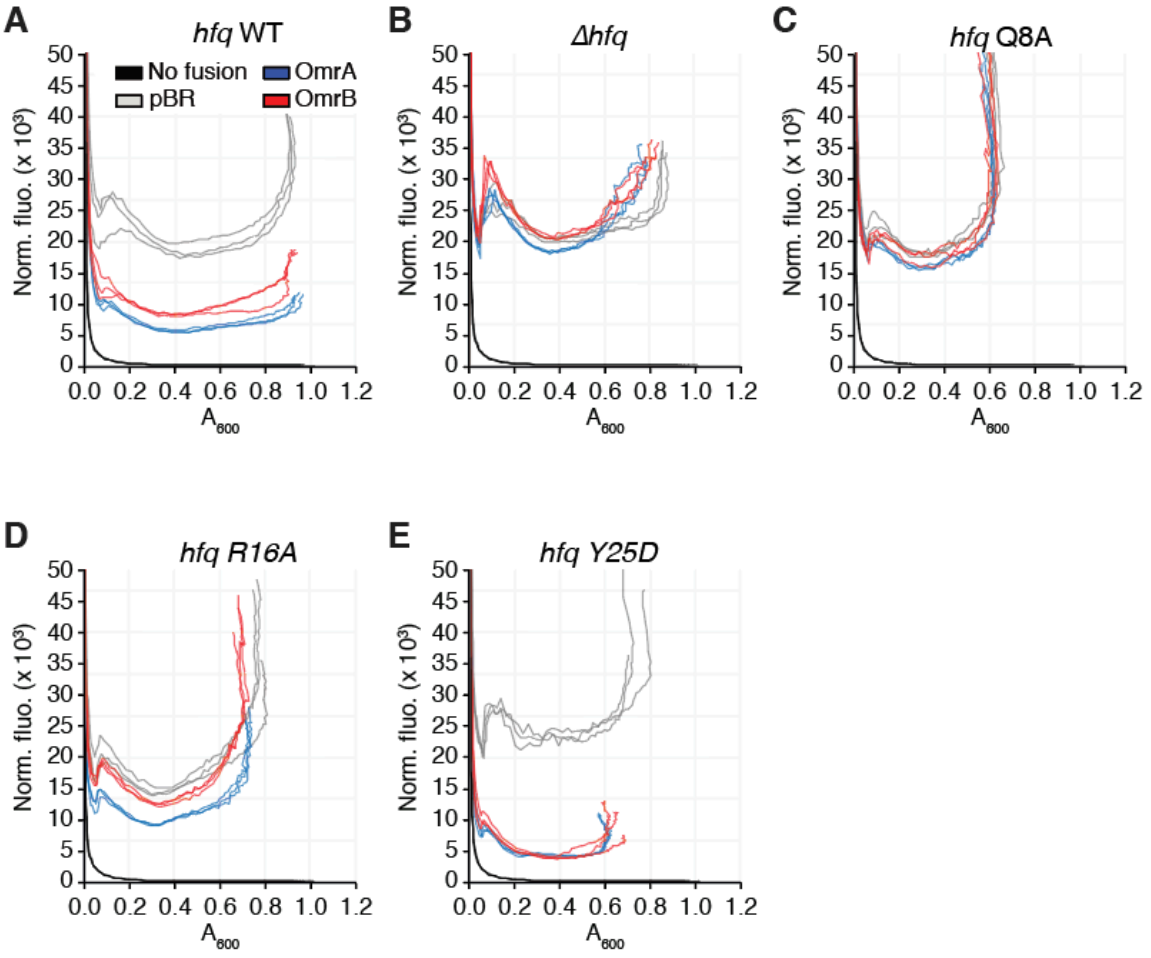

**Supplementary Figure S5. Fluorescence assays using the OmpR$_{717}$-mScarlet fusion.** **(A-E)** Experiments were performed when overexpressing OmrA, OmrB or when using the empty vector (pBR). A control was performed without the fluorescent construct (no fusion). Experiments were performed in the WT strain (*hfq* WT) **(A)**, Δ*hfq* **(B)**, *hfq* Q8A **(C)**, *hfq* R16A **(D)** and *hfq* Y25D **(E)**. These raw data were used to calculate the relative FPA (fluorescence per absorbance) shown in Figure 4C.

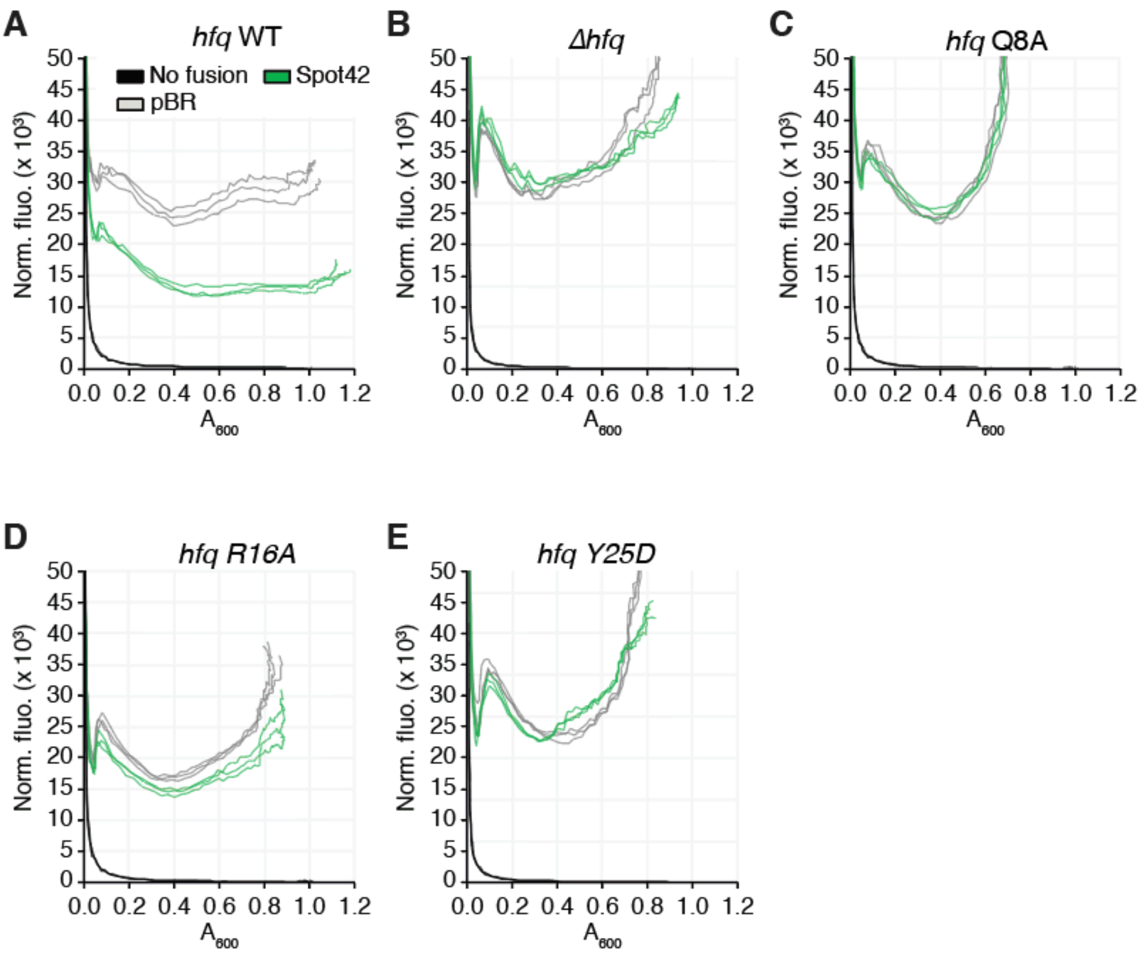

**Supplementary Figure S6. Fluorescence assays using the $SdhC_{39}$-mScarlet fusion. (A-E)** Experiments were performed when overexpressing Spot42 or when using the empty vector (pBR). A control was performed without the fluorescent construct (no fusion). Experiments were performed in the WT strain (*hfq* WT) **(A)**, Δ*hfq* **(B)**, *hfq* Q8A **(C)**, *hfq* R16A **(D)** and *hfq* Y25D **(E)**. These raw data were used to calculate the relative FPA (fluorescence per absorbance) shown in Figure 4D.

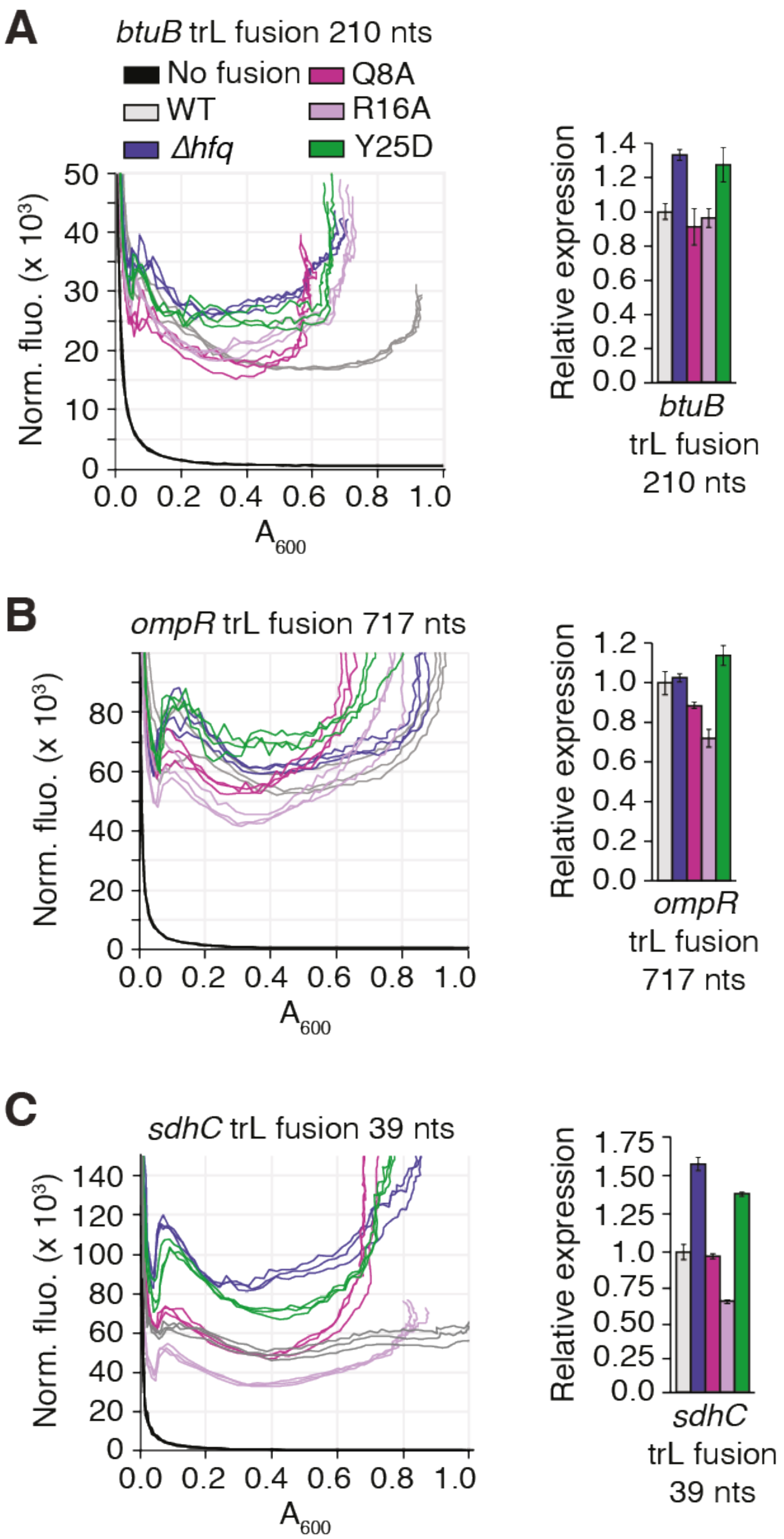

**Supplementary Figure S7. The fluorescence of BtuB$_{210}$-mScarlet (A), OmpR$_{717}$-mScarlet (B) and SdhC$_{39}$-mScarlet (C) fusions was measured in different *hfq***

**backgrounds, in strains transformed with the pBRplac empty vector.** Data are from the datasets shown in Supplementary Figure S4 (*btuB*, A), S5 (*ompR*, B) and S6 (*sdhC*, C). The corresponding relative expression diagrams (right panels) were prepared as in supplementary Figure S2.

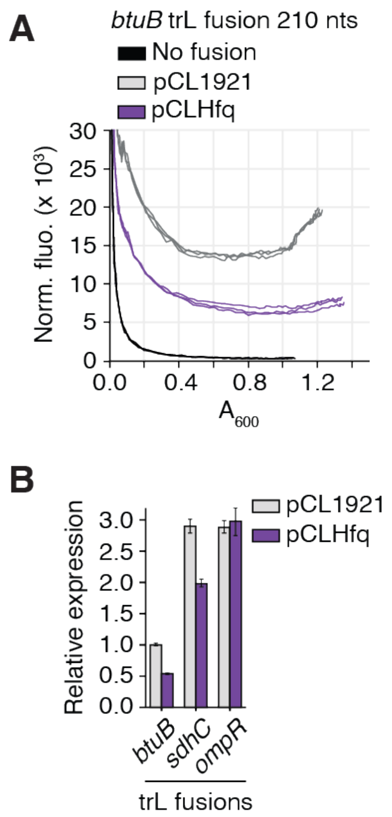

**Supplementary Figure S8. Hfq overproduction represses *btuB* and *sdhC* expression.**
**(A)** Graphic representation is as in Supplementary Figure S2. **(B)** The fluorescence of the $BtuB_{210}$-, $OmpR_{717}$- and $SdhC_{39}$-mScarlet was measured in cells transformed with a plasmid overproducing Hfq (pCLHfq) or the corresponding empty vector (pCL1921) in triplicates.

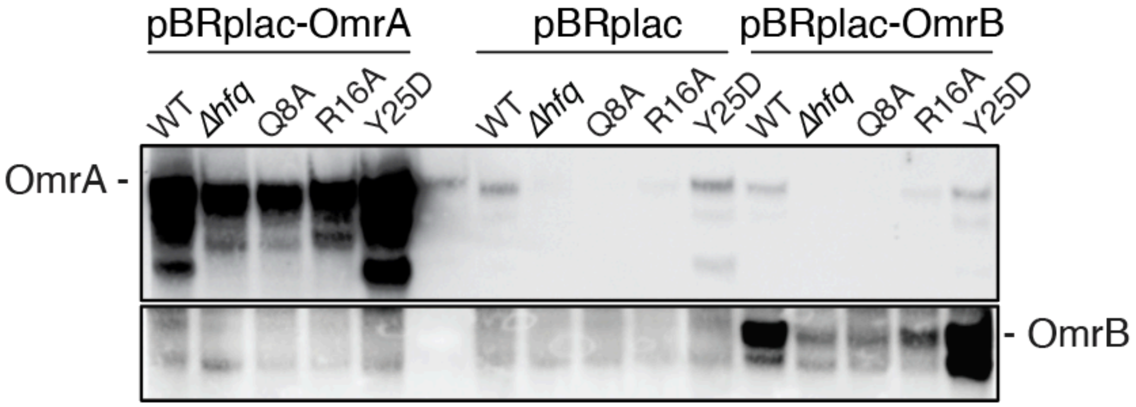

**Supplementary Figure S9. Northern blot analysis of levels of OmrA and OmrB.** Shown are the same Northern blots as in Figure 4E, with a longer acquisition time, so that the chromosomally-expressed OmrA sRNA can now be detected.

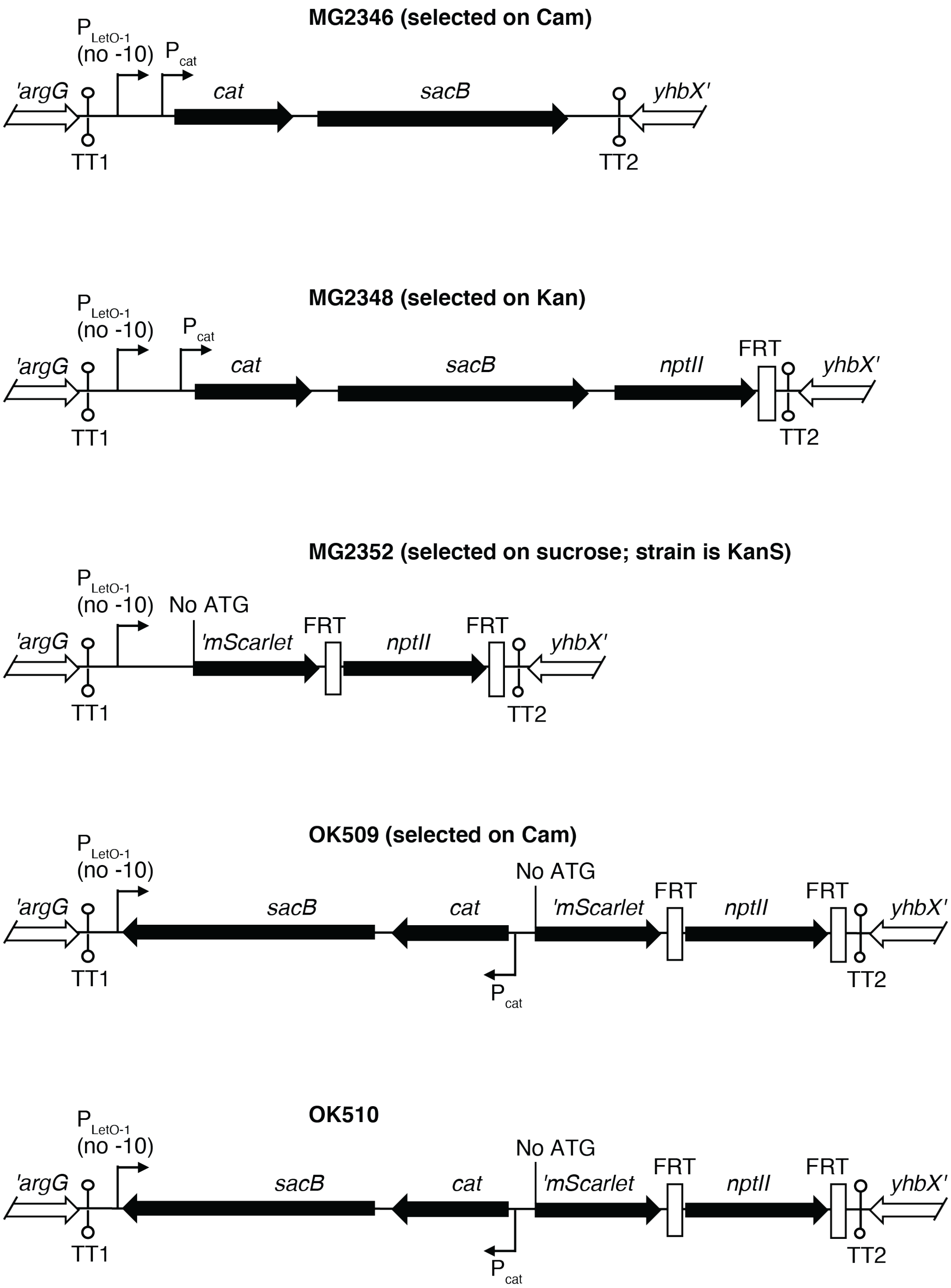

**Supplementary Figure S10. Schematics of the different steps leading to the OK510 strain used for the construction of mScarlet fusions by recombineering.** Shown is the genetic organization of the *argG-yhbX* intergenic region in the different intermediate strains. See supplementary text for details.

### Table S1. Strains and plasmids used in this study.

| Strain name | Characteristics | References |
|---|---|---|
| MG1655 | E. coli reference strain for this study | From F. Blattner's lab |
| DJ480 | MG1655 *ΔlacX74* | D. Jin, NIH |
| DJ624 | MG1655 *ΔlacX74, mal::lacI*$^{q}$ | D. Jin, NIH, used as 'no fusion' control strain in Fig. 4 |
| EM1055 | MG1566 *ΔlacX74* | (17) Same genotype as DJ480, used in Fig. 1 |
| EM1264 | MG1055 *hfq::cat* | (18) Used in Fig. 1 |
| EM1377 | EM1055 *rne131 zce-726::Tn10* | (18) Used in Fig. 1 |
| PM1205 | MG1655 *mal::lacI*$^{q}$ *ΔaraBAD araC+ mini-λ-Tet lacI' ::*$P_{BAD}$*-cat-sacB-lacZ* | (8) |
| BTU1 | PM1205 *lacI′*::$P_{BAD}$-BtuB$_{-240+18}$-LacZ$_{+28}$ | (19) Used in Fig. 2 |
| BTU8 | PM1205 *lacI′*::$P_{BAD}$-BtuB$_{-240+63}$-LacZ$_{+28}$ | This study, recombineering in PM1205, used in Fig. 2 |
| BTU7 | PM1205 *lacI′*::$P_{BAD}$-BtuB$_{-240+81}$-LacZ$_{+28}$ | This study, recombineering in PM1205, used in Fig. 2 |
| BTU20 | PM1205 *lacI′*::$P_{BAD}$-BtuB$_{-240+81}$EP1-LacZ$_{+28}$ | This study, recombineering in PM1205, used in Fig. 2 |
| BTU21 | PM1205 *lacI′*::$P_{BAD}$-BtuB$_{-240+81}$EP2-LacZ$_{+28}$ | This study, recombineering in PM1205, used in Fig. 2 |
| BTU22 | PM1205 *lacI′*::$P_{BAD}$-BtuB$_{-240+81}$EP3-LacZ$_{+28}$ | This study, recombineering in PM1205, used in Fig. 2 |
| BTU6 | PM1205 *lacI′*::$P_{BAD}$-BtuB$_{-240+120}$-LacZ$_{+28}$ | This study, recombineering in PM1205, used in Fig. 2 |
| BTU5 | PM1205 *lacI′*::$P_{BAD}$-BtuB$_{-240+210}$-LacZ$_{+28}$ | This study, recombineering in PM1205, used in Fig. 2 |
| BTU4 | PM1205 *lacI′*::$P_{BAD}$-BtuB$_{-240+420}$-LacZ$_{+28}$ | This study, recombineering in PM1205, used in Fig. 2 |
| BTU2 | PM1205 *lacI′*::$P_{BAD}$-*btuB*$_{-240+18}$-*lacZ*$_{-38}$ | (19) Used in Fig. 2 |
| BTU14 | PM1205 *lacI′*::$P_{BAD}$-*btuB*$_{-240+45}$-*lacZ*$_{-38}$ | This study, recombineering in PM1205, used in Fig. 2 |
| BTU11 | PM1205 *lacI′*::$P_{BAD}$-*btuB*$_{-240+210}$-*lacZ*$_{-38}$ | This study, recombineering in PM1205, used in Fig. 2 |
| BTU10 | PM1205 *lacI′*::$P_{BAD}$-*btuB*$_{-240+420}$-*lacZ*$_{-38}$ | This study, recombineering in PM1205, used in Fig. 2 |
| DJS2604 | MG1655 *Δhfq::cat-sacB, purA::*FRT-*nptII*-FRT | D. Schu, unpublished |
| DJS2609 | MG1655 *hfq WT* | D. Schu, unpublished |
| DJS2927 | MG1655 *Δhfq* | D. Schu, unpublished |
| JJ0015 | MG1508 *mhpR*-$P_{LtetO-1}$-*fepA*$_{-173+45}$-*lacZ*$_{+28}$*Δmini-λ-Tet* | (20) Used in Fig. 3E |
| JJ389 | MG1508 *mhpR*-$P_{LtetO-1}$-*fepA*$_{-146+45}$-*lacZ*$_{+28}$*Δmini-λ-Tet* | Lab collection (strain from J. Jagodnik), used in Fig. S2C |
| JJ416 | MG1508 *mhpR*-$P_{LtetO-1}$-*btuB*$_{-240+99}$-*lacZ*$_{+28}$*Δmini-λ-Tet* | This study; recombineering in MG1508, used in Fig. 3F-G and 5B |
| JJ425 | MG1508 *mhpR*-$P_{LtetO-1}$-*btuBM9*$_{-240+99}$-*lacZ*$_{+28}$ *Δmini-λ-Tet* | This study; recombineering in MG1508, used in Fig. 3F-G |
| JJ426 | JJ416 *ΔomrAB::nptI* | This study; JJ416 + P1 (MG1003), used in Fig. 3B-G |
| JJ427 | JJ425 *ΔomrAB::nptI* | This study; JJ425 + P1 (MG1003), used in Fig. 3B, 3E-G |
| JW3140 | BW25113 *ΔargG ::*FRT-*nptII*-FRT | Keio collection (11) |
| JW5851 | BW25113 *Δpnp::* FRT-*nptII*-FRT | Keio collection (11) |
| KK2560 | MG1655 *hfqQ8A* | D. Schu, unpublished |
| KK2561 | MG1655 *hfqR16A* | D. Schu, unpublished |
| KK2562 | MG1655 *hfqY25D* | D. Schu, unpublished |
| MG1001 | DJ480 *ΔomrA::nptI Δmini-λ-Tet* | This study; recombineering in NM300 |
| MG1002 | DJ480 *ΔomrB::nptI Δmini-λ-Tet* | This study; recombineering in NM300 |
| MG1003 | DJ480 *ΔomrAB::nptI Δmini-λ-Tet* | (10) |

| | | |
|---|---|---|
| MG1432 | DJ624 *mini-λ-Tet* | M. Guillier, unpublished. |
| MG1508 | MG1655 *mal::lacI*$^{q}$*, mini-λ-Tet, mhpR*-P$_{LtetO-1}$-*cat-sacB-lacZ* | (9) |
| MG1582 | MG1508 *mhpR*-P$_{LtetO-1}$-*cirA*$_{-160+30}$-*lacZ*$_{+28}$ *Δmini-λ-Tet* | This study; recombineering in MG1508 |
| MG2288 | MG1582 *ΔomrAB::nptI* | This study; MG1582 + P1 (MG1003), used in Fig. 3E |
| MG2346 | MG1432 *argG*-[TT1-P$_{LtetO-1}$(no -10)-*cat-sacB*-TT2]-*yhbX* | This study |
| MG2348 | MG2346 *argG*-[TT1-P$_{LtetO-1}$(no -10)-*cat-sacB-nptII*-FRT-TT2]-*yhbX* | This study |
| MG2352 | MG2348 *argG*-[TT1-P$_{LtetO-1}$(no -10)-*'mScarlet*(no ATG)-FRT-*nptII*-FRT-TT2]-*yhbX* | This study |
| MG2389 | OK819 *ΔomrAB::nptI* | This study; OK819 + P1 (MG1003), used in Fig. 3C |
| NM300 | DJ480 *mini-λ-Tet* | N. Majdalani, NIH |
| OK509 | MG2352 *argG*-[TT1-P$_{LtetO-1}$(no -10)- *sacB-cat-'mScarlet*(no ATG)-FRT-*nptII*-FRT-TT2]-*yhbX* | This study; recombineering: *cat-sacB* cassette recombined into MG2352 |
| OK510 | MG1432 *argG*-[TT1-P$_{LtetO-1}$(no -10)-*sacB-cat-'mScarlet*(no ATG)-FRT-*nptII*-FRT-TT2]-*yhbX* | This study; MG1432 + P1 (OK509) |
| OK523 | DJ624 *ΔargG::*FRT-*nptII*-FRT | This study; DJ624 + P1 (JW3140) |
| OK528 | OK510 *argG*-[TT1-P$_{LtetO-1}$-*sdhC*$_{-222+39}$-*mScarlet*$_{+4}$-FRT-*nptII*-FRT-TT2]-*yhbX, Δmini-λ-Tet* | This study; recombineering in OK510 |
| OK529 | OK510 *argG*-[TT1-P$_{LtetO-1}$-*ompR*$_{-35+717}$-*mScarlet*$_{+4}$-FRT-*nptII*-FRT-TT2]-*yhbX, Δmini-λ-Tet* | This study; recombineering in OK510 |
| OK530 | OK523 *ΔargG ::*FRT | This study; elimination of *nptII* in OK523 *via* Flp-FRT recombination |
| OK560 | OK528 *argG*-[TT1-P$_{LtetO-1}$-*sdhC*$_{-222+39}$-*mScarlet*$_{+4}$-FRT-TT2]-*yhbX* | This study; elimination of *nptII* in OK528 *via* Flp-FRT recombination |
| OK561 | OK529 *argG*-[TT1-P$_{LtetO-1}$-*ompR*$_{-35+717}$-*mScarlet*$_{+4}$- FRT-TT2]-*yhbX* | This study; elimination of *nptII* in OK529 *via* Flp-FRT recombination |
| OK564 | OK530 *Δhfq::cat-sacB, purA::*FRT-*nptII*-FRT | This study; OK530 + P1 (DJS2604) |
| OK572 | OK510 *argG*-[TT1-P$_{LtetO-1}$-*btuB*$_{-240+210}$-*mScarlet*$_{+4}$-FRT-*nptII*-FRT-TT2]-*yhbX, Δmini-λ-Tet* | This study; recombineering in OK510 |
| OK577 | OK572 *argG*-[TT1-P$_{LtetO-1}$-*btuB*$_{-240+210}$-*mScarlet*$_{+4}$-FRT-TT2]-*yhbX* | This study; elimination of *nptII* in OK572 *via* Flp-FRT recombination |
| OK581 | OK564 *hfq WT* | This study; OK564 + P1 (DJS2609) (purine prototroph selection) |
| OK582 | OK564 *Δhfq* | This study; OK564 + P1 (DJS2927) (purine prototroph selection) |
| OK583 | OK564 *hfqQ8A* | This study; OK564 + P1 (KK2560) (purine prototroph selection) |
| OK584 | OK564 *hfqR16A* | This study; OK564 + P1 (KK2561) (purine prototroph selection) |
| OK585 | OK564 *hfqY25D* | This study; OK564 + P1 (KK2562) (Selecting for SucrR) |
| OK586 | OK581 *argG*-[TT1-P$_{LtetO-1}$-*btuB*$_{-240+210}$-*mScarlet*$_{+4}$-FRT-TT2]-*yhbX* | This study; OK581 + P1 (OK577) (arginine prototroph selection), used in Fig. 4A-B and 4E |
| OK587 | OK582 *argG*-[TT1-P$_{LtetO-1}$-*btuB*$_{-240+210}$-*mScarlet*$_{+4}$-FRT-TT2]-*yhbX* | This study; OK582 + P1 (OK577) (arginine prototroph selection), used in Fig. 4B and 4E |
| OK588 | OK583 *argG*-[TT1-P$_{LtetO-1}$-*btuB*$_{-240+210}$-*mScarlet*$_{+4}$-FRT-TT2]-*yhbX* | This study; OK583 + P1 (OK577) (arginine prototroph selection), used in Fig. 4B and 4E |
| OK589 | OK584 *argG*-[TT1-P$_{LtetO-1}$-*btuB*$_{-240+210}$-*mScarlet*$_{+4}$-FRT-TT2]-*yhbX* | This study; OK584 + P1 (OK577) (arginine prototroph selection), used in Fig. 4B and 4E |
| OK590 | OK585 *argG*-[TT1-P$_{LtetO-1}$-*btuB*$_{-240+210}$-*mScarlet*$_{+4}$-FRT-TT2]-*yhbX* | This study; OK585 + P1 (OK577) (arginine prototroph selection), used in Fig. 4B and 4E |
| OK596 | OK581 *argG*-[TT1-P$_{LtetO-1}$-*sdhC*$_{-222+39}$-*mScarlet*$_{+4}$-FRT- TT2]-*yhbX* | This study; OK581 + P1 (OK560) (arginine prototroph selection), used in Fig. 4D and 4F |

| | | |
|---|---|---|
| OK597 | OK582 *argG*-[TT1-$P_{LtetO-1}$-*sdhC*$_{-222+39}$-*mScarlet*$_{+4}$-FRT- TT2]-*yhbX* | This study; OK582 + P1 (OK560) (arginine prototroph selection), used in Fig. 4D and 4F |
| OK598 | OK583 *argG*-[TT1-$P_{LtetO-1}$-*sdhC*$_{-222+39}$-*mScarlet*$_{+4}$-FRT- TT2]-*yhbX* | This study; OK583 + P1 (OK560) (arginine prototroph selection), used in Fig. 4D and 4F |
| OK599 | OK584 *argG*-[TT1-$P_{LtetO-1}$-*sdhC*$_{-222+39}$-*mScarlet*$_{+4}$-FRT- TT2]-*yhbX* | This study; OK584 + P1 (OK560) (arginine prototroph selection), used in Fig. 4D and 4F |
| OK600 | OK585 *argG*-[TT1-$P_{LtetO-1}$-*sdhC*$_{-222+39}$-*mScarlet*$_{+4}$-FRT- TT2]-*yhbX* | This study; OK585 + P1 (OK560) (arginine prototroph selection), used in Fig. 4D and 4F |
| OK601 | OK581 *argG*-[TT1-$P_{LtetO-1}$-*ompR*$_{-35+717}$-*mScarlet*$_{+4}$- FRT-TT2]-*yhbX* | This study; OK581 + P1 (OK561) (arginine prototroph selection), used in Fig. 4C |
| OK602 | OK582 *argG*-[TT1-$P_{LtetO-1}$-*ompR*$_{-35+717}$-*mScarlet*$_{+4}$- FRT-TT2]-*yhbX* | This study; OK582 + P1 (OK561) (arginine prototroph selection), used in Fig. 4C |
| OK603 | OK583 *argG*-[TT1-$P_{LtetO-1}$-*ompR*$_{-35+717}$-*mScarlet*$_{+4}$- FRT-TT2]-*yhbX* | This study; OK583 + P1 (OK561) (arginine prototroph selection), used in Fig. 4C |
| OK604 | OK584 *argG*-[TT1-$P_{LtetO-1}$-*ompR*$_{-35+717}$-*mScarlet*$_{+4}$- FRT-TT2]-*yhbX* | This study; OK584 + P1 (OK561) (arginine prototroph selection), used in Fig. 4C |
| OK605 | OK585 *argG*-[TT1-$P_{LtetO-1}$-*ompR*$_{-35+717}$-*mScarlet*$_{+4}$- FRT-TT2]-*yhbX* | This study; OK585 + P1 (OK561) (arginine prototroph selection), used in Fig. 4C |
| OK615 | JJ416 *ΔomrA::nptI* | This study; JJ416 + P1 (MG1001), used in Fig. 3F-G |
| OK616 | JJ416 *ΔomrB::nptI* | This study; JJ416 + P1 (MG1002), used in Fig. 3F-G |
| OK617 | JJ425 *ΔomrA::nptI* | This study; JJ425 + P1 (MG1001), used in Fig. 3F-G |
| OK618 | JJ425 *ΔomrB::nptI* | This study; JJ425 + P1 (MG1002), used in Fig. 3F-G |
| OK632 | MG1508 *mhpR*-$P_{LtetO-1}$-*btuB*$_{-240+99}$mutH1-*lacZ*$_{+28}$*Δmini-λ-Tet* | This study; recombineering in MG1508, used in Fig. 5B |
| OK737 | MG1508 *mhpR*-low$P_{LtetO-1}$-*btuB*$_{-240+99}$-*lacZ*$_{+28}$*Δmini-λ-Tet* | This study; recombineering in MG1508, used in Fig. 5B |
| OK819 | MG1508 *mhpR*-$P_{LtetO-1}$-*btuBM11*$_{-240+99}$-*lacZ*$_{+28}$ *Δmini-λ-Tet* | This study; recombineering in MG1508 |

| Plasmid | Characteristics | Use |
|---|---|---|
| pNM12 | AmpR, pBAD24 derivative | (21) pBAD vector control |
| pBAD-OmrA | *omrA* under $P_{BAD}$ control in pNM12 | (10) OmrA overproduction |
| pBAD-OmrB | *omrB* under $P_{BAD}$ control in pNM12 | (10) OmrB overproduction |
| pCP20 | $FLP^{+}$, λ $cI857^{+}$, λ $p_R$ $Rep^{TS}$, $Amp^R$, $Cam^R$ | (12) *nptII* elimination *via* Flp-FRT recombination |
| pBRplac | $Amp^R$, $Tet^R$, $P_{LlacO-1}$ cloned into pBR322 | (10) vector control |
| pOmrA | *omrA* under $P_{LlacO-1}$ control in pBRplac | (10) OmrA overproduction |
| pOmrAM5' | *omrAM5'* under $P_{LlacO-1}$ control in pBRplac<br>M5'= change of nt 3-6 of OmrA from CAGA to GAAC (*a.k.a. omrAmut2* in (10)) | (10) OmrAM5' overproduction |
| pOmrAM9* | *omrAM9** under $P_{LlacO-1}$ control in pBRplac<br>M9 = change of nt 26-36 of OmrA from AUUCGGUACGC to GCGAAAUACCG | This study; OmrAM9* overproduction |
| pOmrAM11* | *omrAM11** under $P_{LlacO-1}$ control in pBRplac<br>M9 = change of nt 20-22 of OmrA from GAG to CUC | This study; OmrAM11* overproduction |
| pOmrAopt | *omrAopt* under $P_{LlacO-1}$ control in pBRplac<br>opt= change of nt 32-41 of OmrA from UACGCUCUUC to GCUG (*a.k.a. omrAmut8* in the lab stock) | This study; OmrAopt overproduction |

| | | |
|---|---|---|
| pOmrAoptM9* | *omrAoptM9** under $P_{LlacO-1}$ control in pBRplac<br>optM9* = change of nt 26-41 of OmrA from AUUCGGUACGCUCUUC to GCGAAACGUG (*a.k.a. omrAmut8+9* in the lab stock, it is the compensatory change to *btuBM9* variant) | This study; OmrAoptM9* overproduction |
| pOmrB | *omrB* under $P_{LlacO-1}$ control in pBRplac | (10) OmrB overproduction |
| pSpot42 | *spf* under $P_{LlacO-1}$ control in pBRplac | (22) Spot 42 overproduction |
| pCL1921 | SpcR/StrR, pSC101 derivative | (23) |
| pCL*hfq*⁺ | *E. coli hfq* under its natural promoter control in pCL1921 | (24) |

FRT is the Flp recombinase target site.
*mini-λ-Tet* is a λ prophage lacking replication and lytic functions. It provides Red functions required for recombination, and resistance to tetracycline (3).
*mScarlet* is a fluorescent reporter (25); the mScarlet-I gene was optimized for codon usage in *E. coli* (6).
*nptI* and *nptII* are kanamycin resistance ORFs originating from Tn903 and Tn5, respectively.
$P_{LtetO-1}$ and $P_{LlacO-1}$ are hybrid tetracycline and IPTG-inducible promoters, respectively (5).

#### Table S2. Oligonucleotides used in this study.

| Name | Sequence | Use |
|---|---|---|
| **Strains construction: $P_{BAD}$-driven *lacZ* fusions** | | |
| LB18 | ACCTGACGCTTTTTATCGCAACTCTCTACTGTTTCTCCATGCCGGTCC TGTGAGTTAATAG | Forward primer for construction of $P_{BAD}$-*btuB-lacZ* transcriptional and translational fusions |
| LB66 | TAACGCCAGGGTTTTCCCAGTCACGACGTTGTAAAACGACCTGTGCC CAAGCGGAAAATG | Reverse primer for BTU8 construction |
| LB67 | TAACGCCAGGGTTTTCCCAGTCACGACGTTGTAAAACGACAGTATCC GGGCTGGTATCC | Reverse primer for BTU7 construction |
| LB75 | TAACGCCAGGGTTTTCCCAGTCACGACGTTGTAAAACGACGCTGCGC GGCTGTTCAAAAC | Reverse primer for BTU6 construction |
| LB76 | TAACGCCAGGGTTTTCCCAGTCACGACGTTGTAAAACGACCGGAAGA CGGCGCAGCA | Reverse primer for BTU5 construction |
| LB77 | TAACGCCAGGGTTTTCCCAGTCACGACGTTGTAAAACGACGGAACCA TAAACAGCGGAGC | Reverse primer for BTU4 construction |
| LB78 | GTGTGATAAAGAAAGTTAAAATGCCGGATCTGCCGTGACGGAACAC GC | Reverse primer for BTU14 construction |
| LB80 | GTGTGATAAAGAAAGTTAAAATGCCGGATCCGGAAGACGGCGCAGC A | Reverse primer for BTU11 construction |
| LB81 | GTGTGATAAAGAAAGTTAAAATGCCGGATCGGAACCATAAACAGCG GAGC | Reverse primer for BTU10 construction |
| Stop 5'-3' | TAACGCCAGGGTTTTCCCAGTCACGACGTTGTAAAACGACCATAGCT GTTTCCTGTGTGATAAAGAAAGTTAAAATGCCGGATC | Reverse primer to include the stop codon sequence followed by *lacZ* translation initiation region and homology region within *lacZ* (construction of transcriptional fusions) |
| LB69 | GGTCTTCATCATGCCGTAATATTGATG | Forward primer for EP1 mutant fusion construction |
| LB70 | CATCAATATTACGGCATGATGAAGACC | Reverse primer for EP1 mutant fusion construction |
| LB71 | GATGAAACCTGCCCGCATCCTTCTTCTATTG | Forward primer for EP2 mutant fusion construction |
| LB72 | CAATAGAAGAAGGATGCGGGCAGGTTTCATC | Reverse primer for EP2 mutant fusion construction |
| LB73 | CTTCTATTGTGGATCGAATACAATGATTAAAAAAGCTTCGC | Forward primer for EP3 mutant fusion construction |
| LB74 | GCGAAGCTTTTTTAATCATTGTATTCGATCCACAATAGAAG | Reverse primer for EP3 mutant fusion construction |
| **Strains construction: $P_{LtetO-1}$-driven *lacZ* fusions** | | |
| 5'Ptet-cirA | GATAGAGATTGACATCCCTATCAGTGATAGAGATACTGAGCACAATC AAAAAAGGCTGACAAATCAG | $P_{LtetO-1}$-*cirA*$_{-160+30}$-*lacZ*$_{+28}$ construction |
| 3'cirA-lacZ | TAACGCCAGGGTTTTCCCAGTCACGACGTTGTAAAACGACGACCCGT ACGAAAGGGTTCAAC | |
| 5'PtetBtuB-240 | CCCTATCAGTGATAGAGATACTGAGCACGCCGGTCCTGTGAGTTAAT AGGGAATCC | $P_{LtetO-1}$-*btuB*$_{-240+210}$-*mScarlet*$_{+4}$ (or $P_{LtetO-1}$-*btuB*$_{-240+99}$-*lacZ*$_{+28}$) construction |
| 3'lacZ-btuB+99 | CCCAGTCACGACGTTGTAAAACGACGTTAGCAGTAACGACGAGAGTA TCCG | $P_{LtetO-1}$-*btuB*$_{-240+99}$-*lacZ*$_{+28}$ construction |
| btuBmut9rev | CCCAGTCACGACGTTGTAAAACGACGTTAGCAGTAACGACGAGAGTG CGAAACGTGGTATCCTGTGCCCAAGCGG | $P_{LtetO-1}$-*btuBM9*$_{-240+99}$-*lacZ*$_{+28}$ construction |
| AK675 | GCCCGGATACTGAGGTCGTTACTGC | $P_{LtetO-1}$-*btuBM11*$_{-240+99}$-*lacZ*$_{+28}$ construction |
| AK462 | GTGGATGCTTTACAATGACCACCACCGCTTCGCTGC | $P_{LtetO-1}$-*btuB*$_{-240+99}$mutH1-*lacZ*$_{+28}$ construction |
| AK283 | TGACACCATCGAATGGCGCTCCCTATCAGTGATAGAGATGGACATCC CTATC | low$P_{LtetO-1}$-*btuB*$_{-240+99}$-*lacZ*$_{+28}$ construction (amplification using JJ416 strain as template) |
| Ptet-55-12For | CTCCCTATCAGTGATAGAGATTGACATCCCTATCAGTGATAGAG | Used to increase the length of the homology region with $P_{LtetO-1}$ for recombineering if needed (used also for the construction of the mScarlet fusions) |
| lacZ28-66rev | AACGCCAGGGTTTTCCCAGTCACGACGTTGTAAAACGAC | Used to increase the length of the homology region with *lacZ* for recombineering if needed |

| | | |
|---|---|---|
| **Strains construction : $P_{LtetO-1}$-driven *mScarlet* fusions** | | |
| Ptetno-10-cat-for | CACATGACGCGCTAGCTCCCTATCAGTGATAGAGATTGACATCCCTATCAGTGATAGAGGCCAGGAATAGCCAGGAATTAAAATGAGACGTTG | Amplification of the [TT1-$P_{LtetO-1}$(no -10)-*cat-sacB*-TT2] cassette for construction of the MG2346 strain Amplification of *nptII*-FRT cassette for construction of the MG2348 strain |
| Ter-catsacRev | GATGTAAGGTTGAAAAATAAAAACGGCGCTAAAAAGCGCCGTTTTTTTTGACGGTGGTAATCAAAGGGAAAACTGTCCATATG | |
| Ter-Ptet-for | GGTACCAAATTCCAGAAAAGAGGCCTCCCGAAAGGGGGGCCTTTTTTCGTTTTGGTCCAACTAGTCCGATGCGACACACATGACGCGCTAGCTCCCTATC | |
| yhbX-Ter-rev | ATAAGCAAGGTAACCCCACCCCTGAAGGGCAGGGTTGATGTAAGGTTGAAAAATAAAAACGGCGCTAAA | |
| argG-Ter-for | GCCGCAGGTGGAGAATCTGGAAAACAAAGGCCAGTAATTCGCCTCGGTACCAAATTCCAGAAAAGAGGCCTCCCGAAAG | |
| sacB-KanR-For | CCTTGAACAAGGACAATTAACAGTTAACAAATAAAAACGCGAGGATCGTTTCGCATGATTGAACAAG | Amplification of *nptII*-FRT cassette for construction of the MG2348 strain |
| FRT-sacB-Rev | CAAAGGGAAAACTGTCCATATGCACAGATGAAAACGGTGTAGTATGAATATCCTCCTTAGTTCCTATTCC | |
| Ptetno-10-mSC-For | GAGATTGACATCCCTATCAGTGATAGAGGCCAGGAATACATTCGGGCGTGAGCAAAGGCGAAG | Amplification of 'mScarlet-FRT cassette (with primer Ptet-55-12For) for construction of the MG2352 strain |
| mSc-FRT-Rev | GCGAAACGATCCTCATCCTGTCGAAGTTCCTATTCTCTAGAAAGTATAGGAACTTCATTATTTATACAGTTCGTCCATGCC | |
| postFRT-KanR-Rev | GAACCTGCGTGCAATCCATCTTGTTCAATCATGCGAAACGATCCTCATCCTGTC | |
| AK411 | ATCAGTGATAGAGATTGACATCCCTATCAGTGATAGAGGCATTTGAGAAGCACACGG | *cat-sacB* amplification for construction of OK509 strain |
| AK412 | ATTCTTTTATTACTGCTTCGCCTTTGCTCACGCCCGAATGAGACGTTGATCGGCAC | |
| AK420 | AACGCATAAATTCTTTTATTACTGCTTCGCCTTTGCTCAC | $P_{LtetO-1}$-*btuB*$_{-240+210}$-*mScarlet*$_{+4}$ construction (with primers 5'PtetBtuB-240 and Ptet-55-12For) |
| AK451 | CTGCTTCGCCTTTGCTCACCGGAAGACGGCGCAGCACATC | |
| 5'PtetompR-35+30-lacZ | TAGAGATTGACATCCCTATCAGTGATAGAGATACTGAGCACTTACAAATTGTTGCGAACCTTTGGGAGT | $P_{LtetO-1}$*ompR*$_{-35+717}$-*mScarlet* $_{+4}$ construction |
| ompR+717-mScrev | GAAACGCATAAATTCTTTTATTACTGCTTCGCCTTTGCTCACTGCTTTAGAGCCGTCCGGTACAAAG | |
| Ptet-sdhC-222for | GACATCCCTATCAGTGATAGAGATACTGAGCACAGGTCTCCGGAACACCCTGCAATC | $P_{LtetO-1}$*sdhC*$_{-222+39}$-*mScarlet* $_{+4}$ construction |
| sdhC+39-mScrev | GAAACGCATAAATTCTTTTATTACTGCTTCGCCTTTGCTCACCAGATTAACAGGTCTTTGTTTTTTCAC | |
| **Strains construction : *omrA/B* deletions** | | |
| OmrA-kan3 | CGCGAGCGACAGTAAATTAGGTGCGAAAAAAAACCTGCGCATCCGCGCAGGTTAGAAAAACTCATCGAGCA | a.k.a. ΔomrABrev in (10); *ΔomrA::nptI* construction |
| OmrA-kan5 | CCCTTCATTCCTTTGCGTTTTCTCGCTGGCGAAGAGTCGTCGTGCAGACAAAGCCACGTTGTGTCTCAA | *ΔomrA::nptI* construction |
| OmrB-kan3 | CGCGAGCGACAGTAAATTAGGTGCGAAAAAAAACCTGCGCATCCGCGCAGGTTAGAAAAACTCATCGAGCA | *ΔomrB::nptI* construction |
| OmrB-kan5 | GCGAAACGCTGTTGCGATTGACCGCTGGTGGCGTTTGGCTTCAGGTTGCAAAGCCACGTTGTGTCTCAA | a.k.a. ΔomrABfor in (10); *ΔomrB::nptI* construction |
| **Verification/sequencing** | | |
| AK87 | CCTTCTCCTGCTCTCCCTTAAGCG | *argG* check |
| AK88 | CTCACGGGTTGTGGATGCAAAC | *argG* check |
| AK387 | CTTCACCTTCACCCTCGATTTC | *mScarlet* fusions check |
| AK418 | ACTGGCCTGCTTTCCTCCTC | *mScarlet* fusions check |
| AK430 | CGTTCAGGAACGGATCTGC | *hfqQ8A* verification |
| AK431 | TGGAACACGTTCCCGAGC | *hfqR16A* verification |
| AK432 | GCTTAATACCATTCACCAAATC | *hfqY25D* verification |
| antiHfqout | GATGTGTACCAGTACCGCCT | *hfq* check (E. Hajnsdorf) |
| lacZ96-120(-) | GCTATTACGCCAGCTGGCGAAAGGG | *lac* fusions check |
| mHfqout | CAGGCGCGTGACGAAGTATT | *hfq* check (E. Hajnsdorf) |
| mhpR848-872 | ATCTTCCGGCGCTACAACGGGTAGC | *lac* fusions check |
| seqOmrArev | GCATTCCAACTCCCTTTGCTC | *omrAB* check |
| seqOmrBfor | GGTGGCGTGTTTTCATCGTGG | *omrAB* check |
| **Plasmids construction** | | |

| | | |
|---|---|---|
| OmrAmut8for | GATTGGTGAGATTATTCGGGCTGGTACCCTGTCTCTTGCACC | Constructing pOmrAopt |
| OmrAmut8rev | GGTGCAAGAGACAGGGTACCAGCCCGAATAATCTCACCAATC | |
| OmrAmut9for | GGTATTGATTGGTGAGATTGCGAAATACCGTCTTCGTACCCTGTCTCTTG | Constructing pOmrAM9* |
| OmrAmut9rev | CAAGAGACAGGGTACGAAGACGGTATTTCGCAATCTCACCAATCAATACC | |
| OmrAmut8+9for | GATTGGTGAGATTGCGAAACGTGGTACCCTGTCTCTTGCACC | Constructing pOmrAoptM9* |
| OmrAmut8+9rev | GGTGCAAGAGACAGGGTACCACGTTTCGCAATCTCACCAATC | |
| OmrAmut11for | CCCAGAGGTATTGATTGGTCTCATTATTCGGTACGCTCTTCGTACCC | Constructing pOmrAoptM11* |
| OmrAmut11rev | GAAGAGCGTACCGAATAATGAGACCAATCAATACCTCTGGGGACGTC | |
| **Templates for T7 *in vitro* transcription and toeprint** | | |
| T7btuBfor | GGTAATACGACTCACTATAGCGCCAACGTCGCATCTGGTTCTC | For *in vitro* transcription of *btuB* (-80+161) region |
| btuBrev | ATATCCTGACGGGTCACAACGG | |
| T7OmrAfor | TAATACGACTCACTATAGGGCCCAGAGGTATTGATTGGTGAG | For *in vitro* transcription of *omrA* |
| OmrArev | AAAAAAAACCTGCGCATCCGCGC | |
| 3995JG | GTGCCCAAGCGGAAAATGCC | Fig. 5E |
| **Templates for T7 *in vitro* transcription of radiolabeled probes for Northern blots** | | |
| T7btuBprobe-for | TAATACGACTCACTATAGCGTACGCCATCAATTAACACCAAC | *in vitro* transcription of the antisense of *btuB* (+85+312) mRNA |
| btuBprobe-rev | GTCGTTACTGCTAACCGTTTTGAACAG | |
| T7OmrAprobe-for | TAATACGACTCACTATAGGGAAAAAAAACCTGCGCATCCGCGC | *in vitro* transcription of the antisense of OmrA sRNA |
| OmrAprobe-rev | CCCAGAGGTATTGATTGGTGAG | |
| T7OmrBprobe-for | TAATACGACTCACTATAGGGAAAAAAAACCTGCGCATCTGCGC | *in vitro* transcription of the antisense of OmrB sRNA |
| OmrBprobe-rev | CCCAGAGGTATTGATAGGTGAAGTC | |
| EM192 | TAATACGACTCACTATAGGGAGATGCCTGGCAGTTCCCTACTC | *in vitro* transcription of the antisense of 5S rRNA |
| EM193 | TGCCTGGCGGCAGTAGCG | |
| EM293 | TAATACGACTCACTATAGGGAGACGCTTTACGCCCAGTAATTCC | *in vitro* transcription of the antisense of a 16S rRNA region |
| EM294 | CTCCTACGGGAGGCAGCAGT | |
| **Biotinylated probes for Northern blots** | | |
| 3'OmrA-probe | BioTEG-CTGCGCATCCGCGCAGGTTGGTGCAAGAGACAGGGTAC | Fig. 3D, S3C |
| OmrA-probe | BioTEG-CAGGTTGGTGCAAGAGACAGGGTACGAAGAGCGTACCG | Fig. 3F, 4E, S2C and S9 |
| OmrB-probe | BioTEG-CGCAGGCTGGTGTAATTCATGTGCTCAACCCGAAGTTGA | Fig. 3F, 4E and S9 |
| Spot42-probe | BioTEG-GTAAAAGGTCTGAAAGATAGAACATCTTACCTCTGTACCC | Fig. 4F |
| SsrA-probe | BioTEG-CGCCACTAACAAACTAGCCTGATTAAGTTTTAACGCTTCA | Fig. 3F, 4F |
| 5S-probe | BioTEG-CTACCATCGGCGCTACGGCGTTTCACTTCTGAGTTCG | Fig. 4E |
